# Prediction of protein biophysical traits from limited data: a case study on nanobody thermostability through NanoMelt

**DOI:** 10.1101/2024.09.13.612907

**Authors:** Aubin Ramon, Mingyang Ni, Olga Predeina, Rebecca Gaffey, Patrick Kunz, Shimobi Onuoha, Pietro Sormanni

## Abstract

1

In-silico prediction of protein biophysical traits is often hindered by the limited availability of experimental data and their heterogeneity. Training on limited data can lead to overfitting and poor generalisability to sequences distant from those in the training set. Additionally, inadequate use of scarce and disparate data can introduce biases during evaluation, leading to unreliable model performances being reported. Here, we present a comprehensive study exploring various approaches for protein fitness prediction from limited data, leveraging pre-trained embeddings, repeated stratified nested cross-validation, and ensemble learning to ensure an unbiased assessment of the performances. We applied our framework to introduce NanoMelt, a predictor of nanobody thermostability trained with a dataset of 640 measurements of apparent melting temperature, obtained by integrating data from the literature with 129 new measurements from this study. We find that an ensemble model stacking multiple regression using diverse sequence embeddings achieves state-of-the-art accuracy in predicting nanobody thermostability. We further demonstrate NanoMelt’s potential to streamline nanobody development by guiding the selection of highly stable nanobodies. We make the curated dataset of nanobody thermostability freely available and NanoMelt accessible as a downloadable software and webserver.

**Significance Statement:** Rapidly predicting protein biophysical traits with accuracy is a key goal in protein engineering, yet efforts to develop reliable predictors are often hindered by limited and disparate experimental measurements. We introduce a framework to predict biophysical traits using few training data, leveraging diverse machine learning approaches via a semi-supervised framework combined with ensemble learning. We applied this framework to develop NanoMelt, a tool to predict nanobody thermostability trained on a new dataset of apparent melting temperatures. Nanobodies are increasingly important in research and therapeutics due to their ease of production and small size, which allows deeper tissue penetration and seamless combination into multi-specific compounds. NanoMelt outperforms available methods for protein thermostability prediction and can streamline nanobody development by guiding the design and selection of highly stable nanobodies during discovery and optimization campaigns.

## 3 Introduction

Protein fitness, defined as the ability of a protein to perform its biological function, encompasses various functional and biophysical properties, including binding affinity, enzymatic activity, solubility, and conformational stability. In silico prediction and optimization of protein fitness have become increasingly prevalent, as they reduce the need for time- and resource-intensive experiments (1, 2). Coarse but rather reliable predictions of protein fitness can now be made ab initio using property-agnostic models. For instance, large language models like ESM (3, 4) can assign probabilities to whether a polypeptide sequence is protein-like and capable of folding and functioning, while models like AlphaFold (5) can predict native protein states, potentially inferring function.

However, the quantitative prediction of biophysical traits that underpin fitness, such as enzymatic activity, binding affinity, or native-state stability, requires property-specific approaches. Energy-based or heuristics ab initio calculations remain challenging, as they need high-resolution structural information as input, and rely on computationally demanding calculations, or on rather inaccurate approximations, which hamper many applications (6–8). Therefore, accurate, rapid, practically useful predictions with the potential to replace experiments generally require rather large amounts of diverse protein-specific or family-specific experimental data for the training of data-driven approaches.

A key limitation in this field is the scarcity of diverse experimental data, which hinders model generalisability to novel or distant sequences (9). Moreover, the available data are often heterogeneous and poorly consistent, as measurements are frequently performed using different techniques or under varying conditions. When such inconsistent data are aggregated, it further complicates the training of effective predictors, as the variability introduced by differing methodologies can obscure true biophysical signals and reduce model accuracy. This necessitates robust training and performance assessment procedures to optimize data use and prevent biases, which are crucial when dealing with limited data (10, 11).

Various strategies have been employed to improve predictions in the context of limited data (12). Transfer learning, for instance, leverages protein representations from pre-trained models on large databases for supervised downstream tasks with fewer data (13, 14). Few-shot learning and semi-supervised learning further refine these models with minimal labelled data (15). Ensemble learning, which combines multiple models, enhances robustness and accuracy (16). Techniques such as bagging and stacking reduce overfitting and improve generalisation by integrating the strengths of individual models (17).

In this work, we introduce a novel framework for learning protein biophysical traits from limited and heterogeneous labelled data. We first estimate the data requirements for effective generalisation using unconstrained artificial data, and then apply this framework to develop NanoMelt, a predictor of nanobody thermostability. Thermostability, a crucial fitness attribute, quantifies a protein’s resistance to denaturation upon heating and correlates strongly with conformational stability (18) and resistance to physicochemical stresses (19). It significantly influences protein developability, affecting expression levels (20), colloidal stability, aggregation propensity (21), and pharmacokinetics (22). Typically quantified by the melting temperature (T_m_) (23), thermostability is often determined via techniques such as differential scanning fluorimetry (DSF), differential scanning calorimetry (DSC), and circular dichroism (CD). Despite the existence of large datasets like ProThermDB (24), which contains approximately 31,000 data points, models developed (6, 25–28) from these datasets often exhibit poor predictive performance for protein subclasses with limited representation, such as monoclonal antibodies (mAbs). Fine-tuning on antibody-specific measurements has been necessary to achieve satisfactory predictions in these cases (29).

Nanobodies, or single-domain antibodies, are small antibody fragments that typically exhibit good expressibility, solubility, stability, and a binding affinity comparable to full antibodies (30). Since their discovery (31), nanobodies have found numerous biological applications (32, 33) and garnered significant therapeutic interest (34), especially following the approval of Caplacizumab and the rise of multi-specific and multi-paratopic drugs. However, public-domain data on nanobody thermostability are limited. The NbThermo (35) database, which contains 548 datapoints, is the most recent compilation of nanobody melting temperatures. The heterogeneity in experimental conditions and methodologies across these measurements results in poorly consistent data, complicating the prediction of nanobody thermostability.

In this study, we present a comprehensive exploration of protein fitness learning with limited data, applying our framework to predict nanobody thermostability. We curated a novel dataset of 640 melting temperatures of unique nanobody sequences, by further cleaning and removing redundancies from NbThermo, merging it with other sources, and performing 129 new measurements. We developed NanoMelt by using an ensemble learning strategy that combines various regression techniques and sequence embeddings. Our approach ensures optimal data utilization and robust performance via a repeated stratified nested cross-validation procedure. We further demonstrate NanoMelt’s potential to streamline nanobody development by guiding the selection and design of highly stable nanobodies. We have made the curated nanobody thermostability dataset freely available, and NanoMelt accessible both as downloadable software and user-friendly webserver.

## 4 Results

### 4.1 Preliminary search of minimal fitness dataset size

In this study, we introduce an extensive protein fitness prediction pipeline optimised for the use of limited data during training and performance assessment (**Fig. 1**), which we apply to the prediction of nanobody thermostability. Training a regression model to accurately predict biophysical traits solely from sequence information remains challenging, with performance largely influenced by the number of labelled sequences available and by how such sequences are represented numerically.

**Fig. 1.**
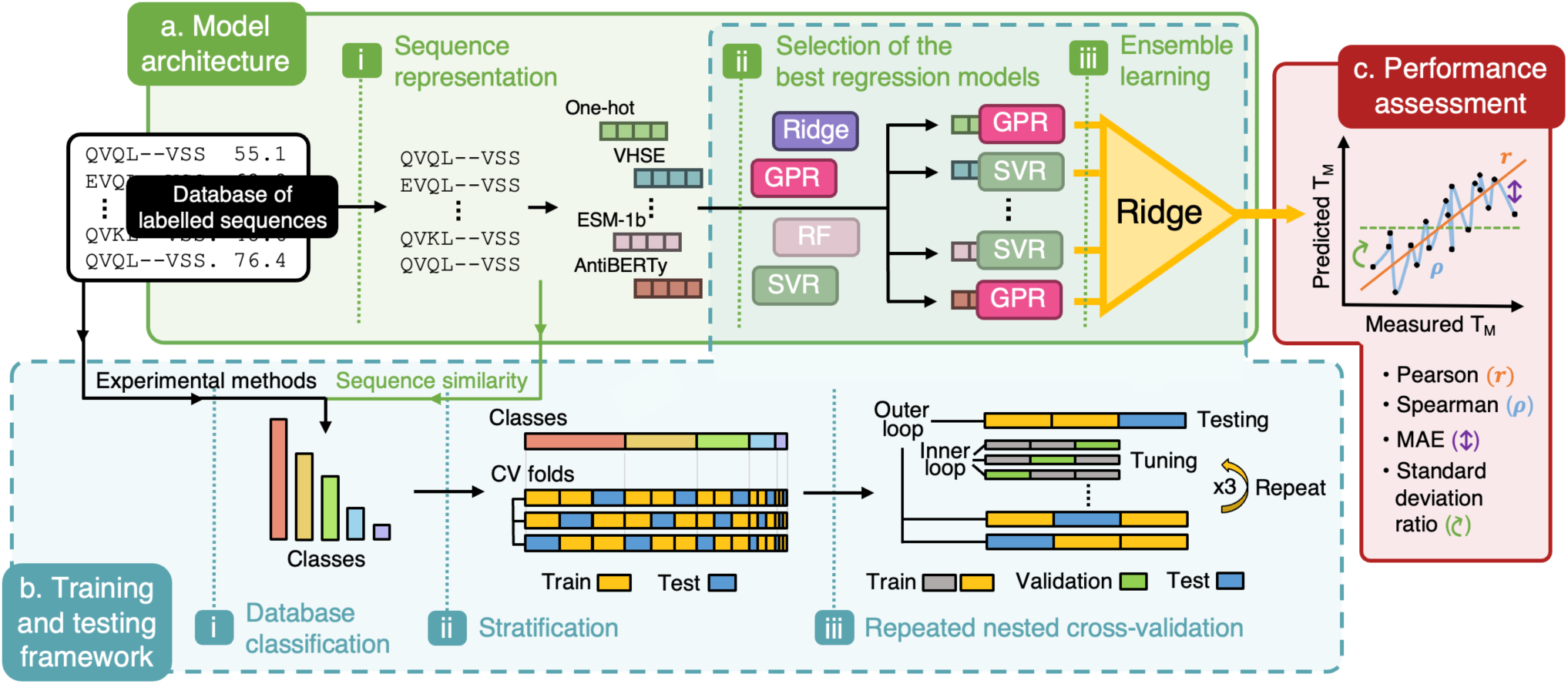
Framework for protein biophysical trait prediction with limited data. **(a)** Model architecture (green): Sequences from a labelled protein database (black box) are represented using multiple embeddings (e.g., ESM-1b, one-hot, VHSE) and processed through various regression models (e.g., ridge, GPR, RF, SVR). The top-performing models for each embedding are combined into a ridge-based ensemble model. **(b)** Training and testing framework (blue): The dataset is clustered by experimental method (e.g., nanoDSF, DSF, DSC, CD in the example of thermostability measurements) and sequence similarity via k-medoids clustering. Stratification ensures consistent class distribution across training (yellow) and testing (blue) splits. Repeated nested cross-validation includes an inner loop for hyperparameter tuning (grey and green) within an outer loop for model testing (yellow and blue), repeated with three random seeds to report averaged performances and their standard deviations. **(c)** Performance assessment (red): Model performance is evaluated using Pearson’s correlation (r), Spearman’s correlation (ρ), mean absolute error (MAE), and standard deviation ratio (SDR), which indicates the model’s tendency to regress towards the dataset mean – a common issue with limited data.

We first aimed to determine the minimum amount of labelled data necessary to train a reliable model capable of generalising to sequences distant from those in the training set, and to evaluate how performance varies with different sequence representations or embeddings. Given the lack of an extensive dataset of nanobody thermostability measurements, we constructed an artificially labelled dataset. The advantage of using artificial data is the ability to generate it in a fully unconstrained manner, enabling the exploration of both large and small datasets of tailored diversity. A total of 10,000 nanobody sequences were selected, and their relative solubility was calculated using the CamSol algorithm (36) (see Methods). CamSol performs complex calculations, combining the physicochemical properties of amino acids to account for both short-range and long-range interactions, ultimately yielding a single solubility score for the whole sequence. We generated our artificial labelled data using two scores: the intrinsic solubility score, derived solely from the amino acid sequence, and the structurally corrected surface score, which also accounts for residue solvent exposure and proximity in the three-dimensional space, as obtained from modelled structures (see Methods). Due to the intricate and non-linear nature of these calculations, we hypothesised that these artificial datasets would approximate reasonably well real datasets of experimentally measured physicochemical properties, at least in terms of prediction complexity.

We trained ridge regression models via repeated cross-validation on datasets of varying sizes, ranging from 20 to 8,000 sequences, and tested all models on the same held-out test set of 2,000 sequences (see Methods and **Fig. S1**). Sequences were represented with eight different embeddings, including categorical (one-hot) and physicochemistry-based (VHSE (37)) encodings of nanobody sequences aligned with the AHo numbering scheme to a fixed length of 149 positions, as well as pre-trained embeddings derived from protein (ESM-1b (38) and ESM-2 (4)), antibody-specific (AbLang (39) and AntiBERTy (40)), and nanobody-specific (nanoBERT (41)) large language models, and from a state-of-the-art nanobody structure predictor (NanobodyBuilder2 (42)), all of which were given non-aligned nanobody sequences as input (see Methods and **Table S1**).

As anticipated, increasing the training set size improved the Spearman’s coefficient (ρ) on the held-out test set and reduced variance across repeated random splits (**Fig. S1**). The categorical one-hot and VHSE encodings exhibited greater overfitting, assessed by the discrepancy between performance on the training and test sets, and showed poorer ability to generalise, requiring a larger number of training sequences to reach a performance plateau on the test set. In contrast, pre-trained embeddings from large language models demonstrated superior generalisation at lower training set sizes. At a training set size exceeding 600, the ESM-1b and ESM-2 begin to reach a performance plateau. For example, at a size of 640, the ESM-1b and ESM-2 show ρ values of 0.871 and 0.888, respectively, compared to 0.605 for the one-hot encoding on the intrinsic solubility score (**Fig. S1a**). Meanwhile, models trained on AntiBERTy, AbLang, and nanoBERT embeddings achieved ρ values of 0.751, 0.793, and 0.802, respectively, suggesting that embeddings trained specifically on antibody or nanobody sequences do not perform as well as those derived from a generic, yet much larger, protein language model. The NanobodyBuilder2 embedding failed to provide sufficient information to the regression model, plateauing at a ρ value of only 0.352 with a training set size of 8,000 sequences. This poor performance is likely due to the NanobodyBuilder2 embedding’s relatively low dimensionality, comprising 128 input features compared to 1,280 for ESM-1b and over 3,000 for one-hot encoding (149*21), as well as its training on a few thousand structures, in contrast to ESM-1b’s 30 million sequences.

In conclusion, consistent with previous reports (14, 16, 43), semi-supervised learning using pre-trained embeddings is the most effective strategy for learning from small protein fitness datasets. Importantly, a training set with more than 600 labelled nanobody sequences appears to provide sufficient information for a sequence-based regression model to yield accurate and generalisable predictions of nanobody biophysical traits. Beyond this threshold, further increases in training data size yield only modest improvements in test-set performance.

### 4.2 Construction of the nanobody thermostability dataset

Apparent melting temperatures (T_m_) of nanobodies have been sporadically reported in the literature. The NbThermo (35) database attempted to compile these thermostability datapoints from various publications. First, we added to the NbThermo a dataset with data from more recent mutational studies, obtaining 511 unique sequences datapoints (see Methods and **Table S2**).

However, our previous analysis on artificial data indicated that these may be insufficient to train a robust model for nanobody fitness prediction. To address this, we generated 129 additional datapoints by characterising nanobodies specifically for this study (**Fig. 2**). These nanobodies included both some already available in our laboratory, and a selection of diverse sequences not previously represented in the database. The largest contribution within NbThermo, comes from Kunz et al. (44), who provided data for 64 diverse nanobodies, all measured under the same conditions, and for which we have the raw fluorescence readouts, meaning that we can re-fit these traces. Therefore, all new measurements were conducted under the same conditions as those used by Kunz et al. (44), and all data were then analysed together, thus yielding a self-consistent dataset of 193 datapoints with uniform technique, protein concentration, buffer, and pH (see Methods). As a control, we reproduced and characterised six nanobodies from Kunz et al., covering a broad range of T_m_ values. The T_m_ values from our measurements (**Fig. S2**) closely align with those from Kunz et al. (44), confirming the high compatibility between the datasets. Measurements were performed at low concentrations to minimise the impact of heat-induced aggregation on equilibrium, which can cause shifts in the measured T_m_ (44) (see Methods and **Fig. 2a**). We carefully extracted the melting temperatures by individually fitting fluorescence emission signals at 350 nm and 330 nm whenever feasible (**Fig. S3**). While fitting the 350/330 nm fluorescence ratio is common practice, this approach can introduce biases under certain conditions (45), as illustrated here by a 3 °C deviation observed in our analysis of the 7KBI melting curves (**Fig. S4**).

**Fig. 2.**
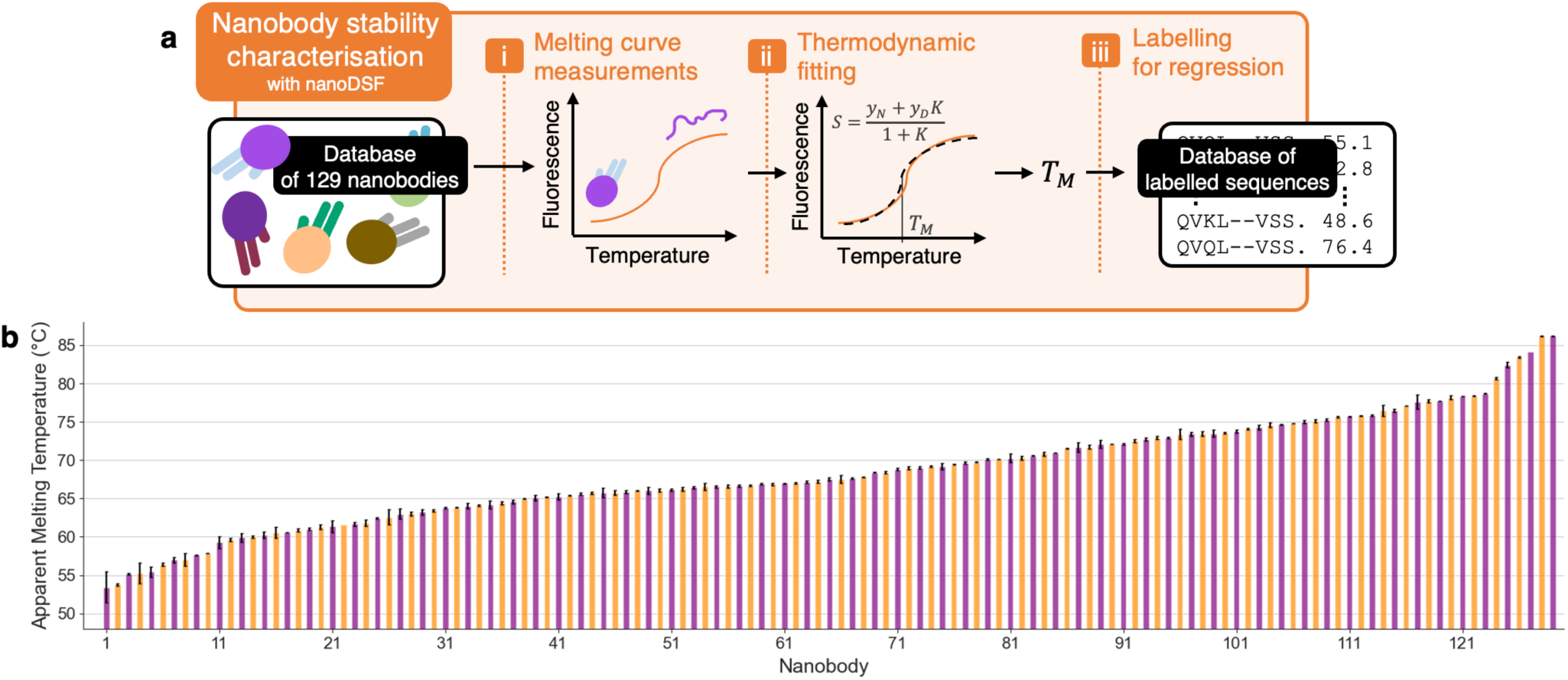
Expansion of nanobody thermostability data. **(a)** Nanobody thermostability characterisation pipeline: The intrinsic fluorescence of 129 nanobodies was measured at increasing temperatures using nanoDSF. The resulting melting curves were fitted to a two-state protein denaturation model to determine the apparent melting temperatures (T_m_) (see Methods). These T_m_ values are then used as labels to train and test our stability predictor, alongside measurements from the literature. **(b)** Fitted T_m_ for the 129 characterised nanobodies: Bar heights are the mean T_m_ from duplicate measurements from separate runs, and error bars the standard deviation. The average error is 0.26 °C, indicating excellent consistency between measurements.

These additional measurements resulted in a final thermostability dataset comprising 640 distinct nanobodies, with T_m_ values ranging from 26.6 °C to 98.2 °C (**Fig. 3a**). The small subset with T_m_ ≤ 42 °C comes from a single study of destabilising mutations (see Methods). The dataset includes measurements from three experimental techniques: 251 from nanoDSF, 205 from circular dichroism (CD), and 165 from sypro-orange DSF (**Fig. 3b**).

**Fig. 3.**
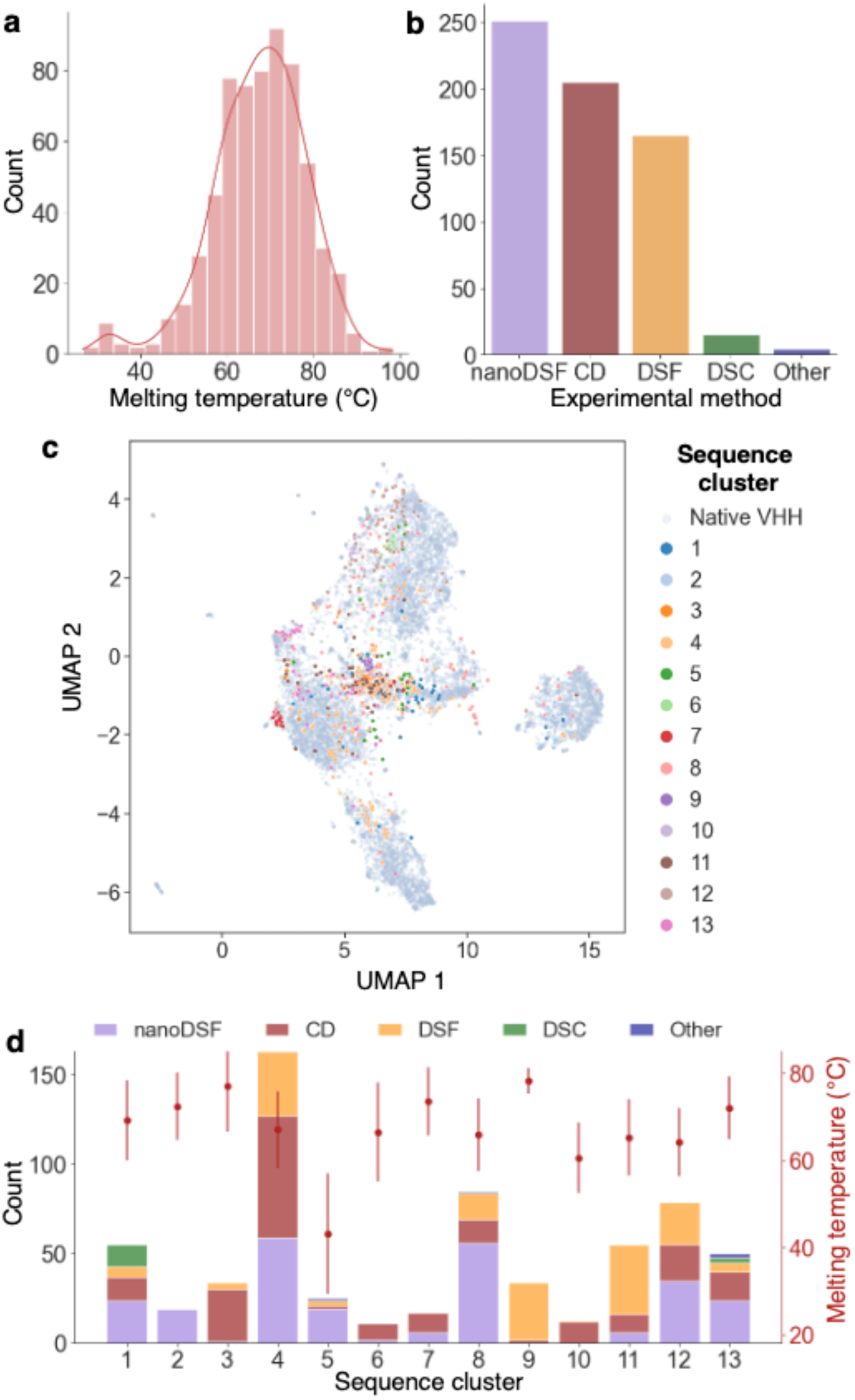
Overview of the Nanobody Thermostability Dataset. **(a)** Histogram showing the distribution of apparent melting temperatures (°C) across the dataset. **(b)** Bar plot depicting the distribution of the experimental methods used. **(c)** UMAP projection (n_neighbors = 20) of the ESM-1b representation of 10,000 native nanobody sequences from the AbNatiV dataset (in grey) overlaid with the 640 sequences from our dataset. Sequences are colour-coded according to their k-medoids cluster number (see Methods). **(d)** Bar plot showing the distribution of experimental methods used within the 13 k-medoids sequence clusters (as coloured in panel c). The average melting temperature and standard deviation for each cluster are indicated by the red points (right y-axis).

We clustered the sequences using the k-medoids algorithm into 13 distinct groups (**Fig. 3c** and Methods). A two-dimensional reduction of these clusters mapped onto the native nanobody space demonstrates the diversity of our dataset, with datapoints roughly evenly distributed across the sequence space (**Fig. 3d**). Additionally, the mutation distance matrix (see Methods and **Fig. S5**) reveals substantial sequence diversity within the dataset, with an average sequence identity as little as 64%, which is rather low for closely related proteins such as nanobodies.

Taken together, these results confirm that we have constructed a broad and diverse dataset of nanobody melting temperatures, achieved also by leveraging significant experimental efforts dedicated to characterising 129 additional nanobodies under consistent conditions to maximise data quality. The complete dataset is available as **Supplementary Dataset 1**.

### 4.3 Selection of regression models

Our analysis using artificial data indicated that 640 datapoints should suffice for training robust models for nanobody fitness prediction. To assess the ability of different regression models to predict nanobody thermostability, we evaluated seven models: three linear (Ridge (46), Huber (47), and Elastic Net (48)) and four non-linear (Random Forest (49) and LightGBM (50), Support Vector Machine (51), and Gaussian Process (52)). Each model used one of eight sequence embeddings as input (**Table S1**) to predict T_m_.

The dataset is limited not only in size but also in consistency, with experimental and analytical variability contributing to differences in apparent melting temperatures. For example, 82 sequences have multiple T_m_ measurements under varying conditions, resulting in a mean pairwise absolute difference of 2.4 °C (**Fig. S6**). To mitigate biases from these inconsistencies and the limited data, we employed a repeated stratified nested cross-validation procedure (11) (**Fig. 1**).

Stratification maintained class distribution across folds, improving generalisability and reliability when training on imbalanced heterogeneous data (53). We stratified based on the experimental method and sequence similarity cluster. Comparison between stratified and random cross-validation showed that stratification improved performance slightly but consistently, and reduced variability (**Fig. S7**). Since diverse experimental methods were used to measure the thermostability in our database, stratification allows to maintain the distribution of these methods in the training data. This ensures that the model learns from a more representative sample, reducing the risk of fitting to the most common experimental methods and compromising the ability to make predictions on the others.

Nested cross-validation allows the efficient use of limited data, while preventing data leakage between training, validation, and test sets, thus keeping the model fine-tuning independent from the model assessment (10). The procedure involves an inner loop for hyperparameter tuning and an outer loop for performance evaluation on an unseen test set, repeated with three random seeds to capture the magnitude of performance variations (**Fig. 1b**).

Results across models and embeddings (**Fig. 4a**, **Tables 1** and **S3**) showed that Gaussian Process Regression (GPR) and Support Vector Regression (SVR) consistently outperformed other models, with higher correlation and lower variance, though with a stronger tendency to overfit (e.g., SVR on ESM-1b: ρ = 0.993 ± 0.004 for training vs. 0.816 ± 0.027 for testing). However, SVR and GPR are expected to fit closely the training data due to their use of kernel functions that model complex data relationships which can capture both the relevant patterns (signal) and the random variations (noise) within the dataset.

**Fig. 4.**
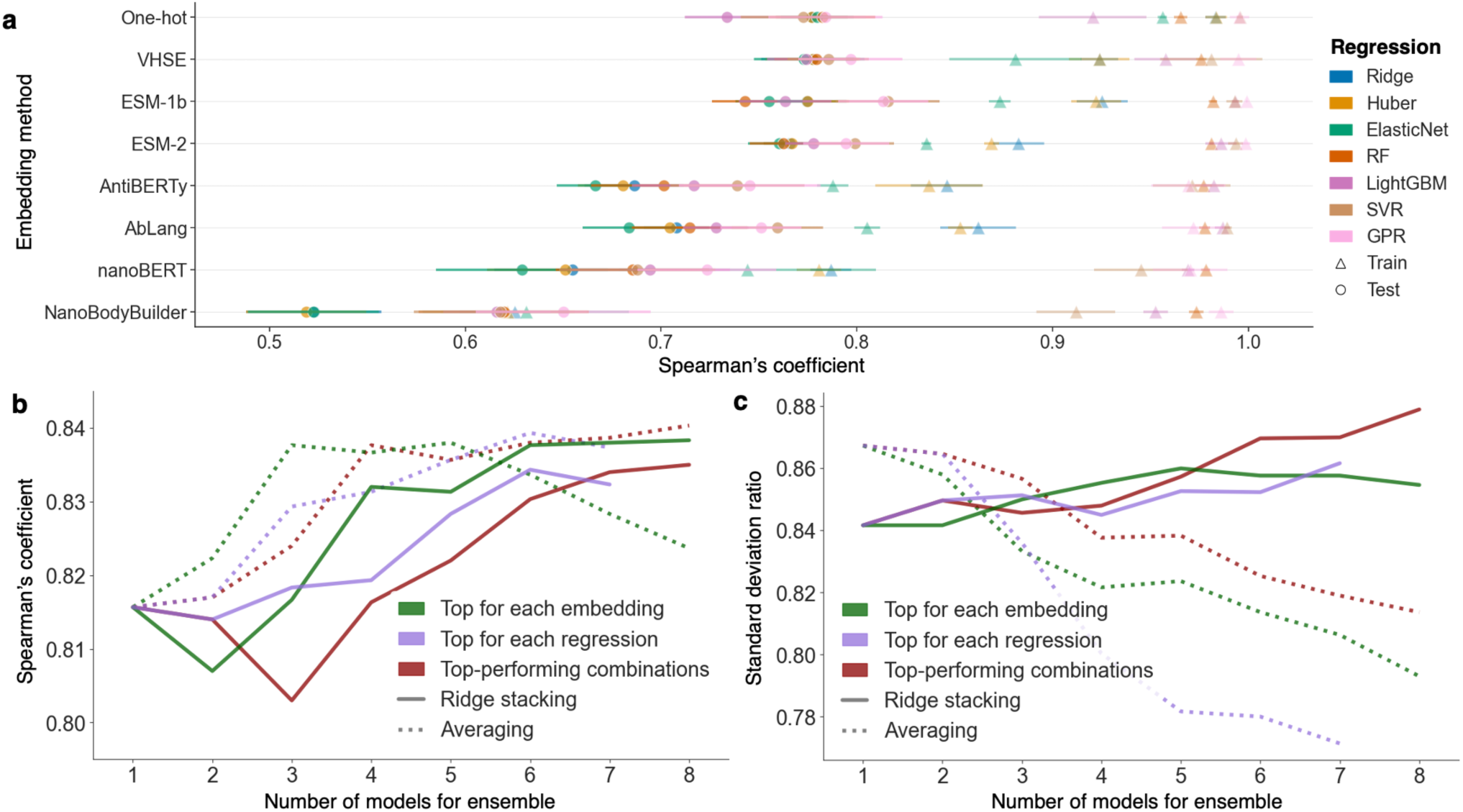
Performance evaluation of regression models for ensemble learning. **(a)** Spearman’s coefficient for selected regression models across different embeddings using repeated nested stratified cross-validation. Each colour represents a regression method (see legend). Triangle markers indicate performance on the training set, while round markers indicate performance on the test set. Error bars denote standard deviations across cross-validation folds and repeats. **(b)** Test-set Spearman’s coefficient and **(c)** standard deviation ratio (SDR) performances for the ridge stacking ensemble (in solid line) and the averaged ensemble (in dotted line) using input predictions from different model selections: top-performing models across diversified embeddings (green, **Table 1**); top-performing embeddings with diversified regression models (purple, **Table S3**); and top-performing combinations of embedding and model from nested stratified cross-validation (red, **Table S3**).

**Table 1.**
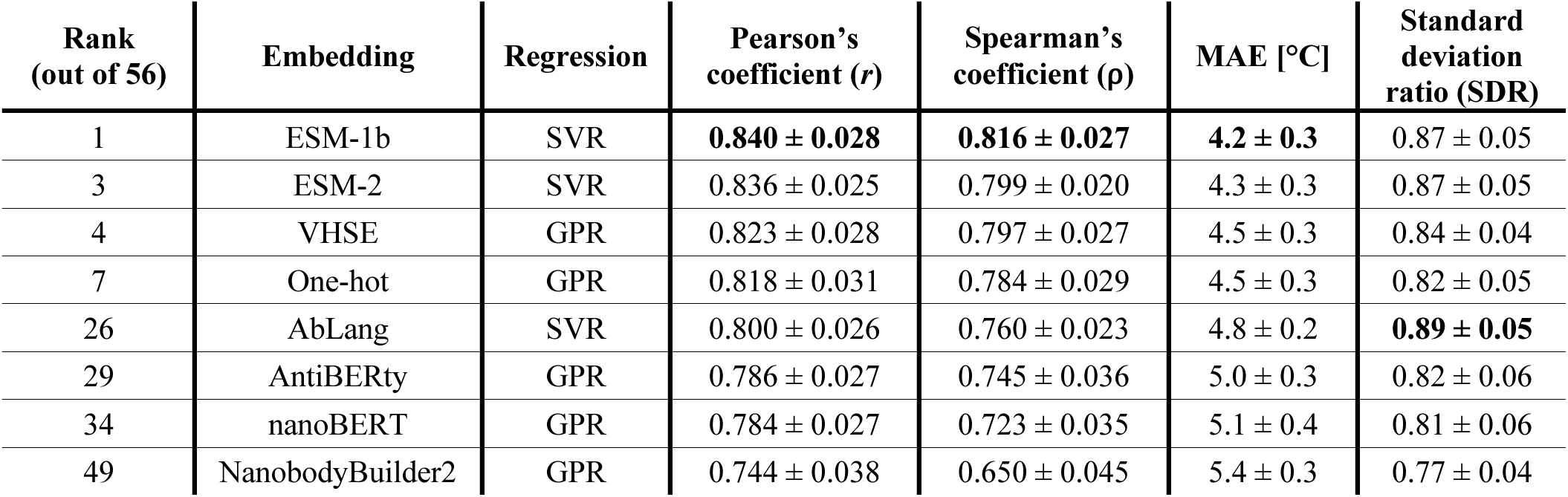
Top-performing regression models for each sequence representation. For each embedding, the table lists the best-performing regression model, ranked according to Spearman’s coefficient. The Pearson’s and Spearman’s coefficients, mean absolute error (MAE), and standard deviation ratio (SDR) are averaged across those on the test folds of the outer loop in the repeated nested cross-validation, and the reported uncertainties are the corresponding standard deviations. Ranks are provided across the 56 possible combinations of regression models and embeddings (see **Table S3**). The best performance in each column is highlighted in bold.

| Rank<br>(out of 56) | Embedding | Regression | Pearson’s<br>coefficient ($r$) | Spearman’s<br>coefficient ($\rho$) | MAE [°C] | Standard<br>deviation<br>ratio (SDR) |
| --- | --- | --- | --- | --- | --- | --- |
| 1 | ESM-1b | SVR | **$0.840 \pm 0.028$** | **$0.816 \pm 0.027$** | **$4.2 \pm 0.3$** | $0.87 \pm 0.05$ |
| 3 | ESM-2 | SVR | $0.836 \pm 0.025$ | $0.799 \pm 0.020$ | $4.3 \pm 0.3$ | $0.87 \pm 0.05$ |
| 4 | VHSE | GPR | $0.823 \pm 0.028$ | $0.797 \pm 0.027$ | $4.5 \pm 0.3$ | $0.84 \pm 0.04$ |
| 7 | One-hot | GPR | $0.818 \pm 0.031$ | $0.784 \pm 0.029$ | $4.5 \pm 0.3$ | $0.82 \pm 0.05$ |
| 26 | AbLang | SVR | $0.800 \pm 0.026$ | $0.760 \pm 0.023$ | $4.8 \pm 0.2$ | **$0.89 \pm 0.05$** |
| 29 | AntiBERTy | GPR | $0.786 \pm 0.027$ | $0.745 \pm 0.036$ | $5.0 \pm 0.3$ | $0.82 \pm 0.06$ |
| 34 | nanoBERT | GPR | $0.784 \pm 0.027$ | $0.723 \pm 0.035$ | $5.1 \pm 0.4$ | $0.81 \pm 0.06$ |
| 49 | NanobodyBuilder2 | GPR | $0.744 \pm 0.038$ | $0.650 \pm 0.045$ | $5.4 \pm 0.3$ | $0.77 \pm 0.04$ |

Among embeddings, the ESM-1b model demonstrated the best overall performance (ρ = 0.816 ± 0.027 with SVR), surpassing those trained on antibody (AntiBERTy: ρ = 0.745 ± 0.036 with SVR) and nanobody (nanoBERT: ρ = 0.723 ± 0.035 with GPR) sequences. VHSE (ρ = 0.797 ± 0.027 with SVR) and one-hot encoding (ρ = 0.784 ± 0.029 with GPR) closely followed but were more prone to overfitting and to regression to the mean (as indicated by lower SDR, **Table 1**). This suggests that pre-trained embeddings offer superior generalisation to sequences distant from the training set.

In summary, repeated stratified nested cross-validation offers a robust framework for comparing models trained on limited, heterogeneous data. Non-linear models with pre-trained embeddings achieved Spearman’s correlations exceeding 0.8 on the test sets.

### 4.4 Ensemble learning

Prediction performance varied significantly depending on the combination of embeddings and regression models used. Ensemble learning, which combines multiple individually trained models into a single predictor, is known to enhance performance by leveraging model diversity, thereby increasing generalisability and robustness when working with limited, heterogeneous data (12, 16).

In this study, predicted test-set T_m_ from individually trained regression models were used as inputs to train the ensemble model. We evaluated different combinations of models for ensemble learning with the same stratified nested cross-validation pipeline (see Methods, **Fig. 4c** and **Table S4**). In addition to the top-ranking combinations (**Table S3**), we tested combinations of the best model for each different embedding (**Table 1**), and of different regression models trained on their best-performing embeddings (**Table S3**). We explored two ensemble strategies for these combinations: averaging the predictions from up to eight models (**Table S3**), and stacking them using a ridge regression meta-model (54) (see Methods). Both approaches achieved comparable Spearman’s correlations (e.g., 0.840 for averaging vs. 0.835 for stacking, using eight top-performing combinations). While averaging performed slightly better when using fewer models (**Fig. 4b**), ridge stacking exhibited a greater resistance to regression towards the mean, as indicated by a higher standard deviation ratio (SDR) (e.g., SDR = 0.81 for averaging vs. 0.88 for stacking, using eight top-performing combinations, **Fig. 4c**). Due to the substantial increase in regression to the mean effect when averaging, ridge regression stacking was selected as the ensemble approach for the remainder of the study.

Performance consistently improved as the number of input predictions increased (**Fig. 4b** and **Table S4**). Notably, the ensemble model based on the top four different embeddings approached the highest Spearman’s correlation (ρ = 0.832), even if the individual models that underpin it have lower performance than the four employed in the corresponding top-performing ridge stacking (**Fig. 4b,c**). Although stacking more than four top-performing models slightly improved the SDR (0.87 for six models vs. 0.86 for four different embeddings), the four-embedding approach was preferred for the remainder of the study due to substantially reduced computational demand at inference time.

Additional strategies to further improve ensemble learning were attempted by incorporating more information besides the predicted input temperatures, following approaches similar to the DeepSTABp framework (26). Attempted augmentations included concatenating the input sequence embeddings or adding experimental conditions as input features (see Methods), but these did not significantly improve model performance (**Table S5**) and were thus not implemented in the final model.

This investigation culminated in an ensemble model that uses regression models trained on four diverse embeddings for nanobody thermostability prediction, which we call NanoMelt. Test-set predictions across the entire dataset of 640 datapoints, through various test folds of the outer cross-validation layer, are shown in **Figure 5**. The model demonstrates robust predictive performance, achieving a Pearson’s correlation of 0.853, a Spearman’s correlation of 0.832, a MAE of 4.1 °C, and an SDR of 0.86.

**Fig. 5.**
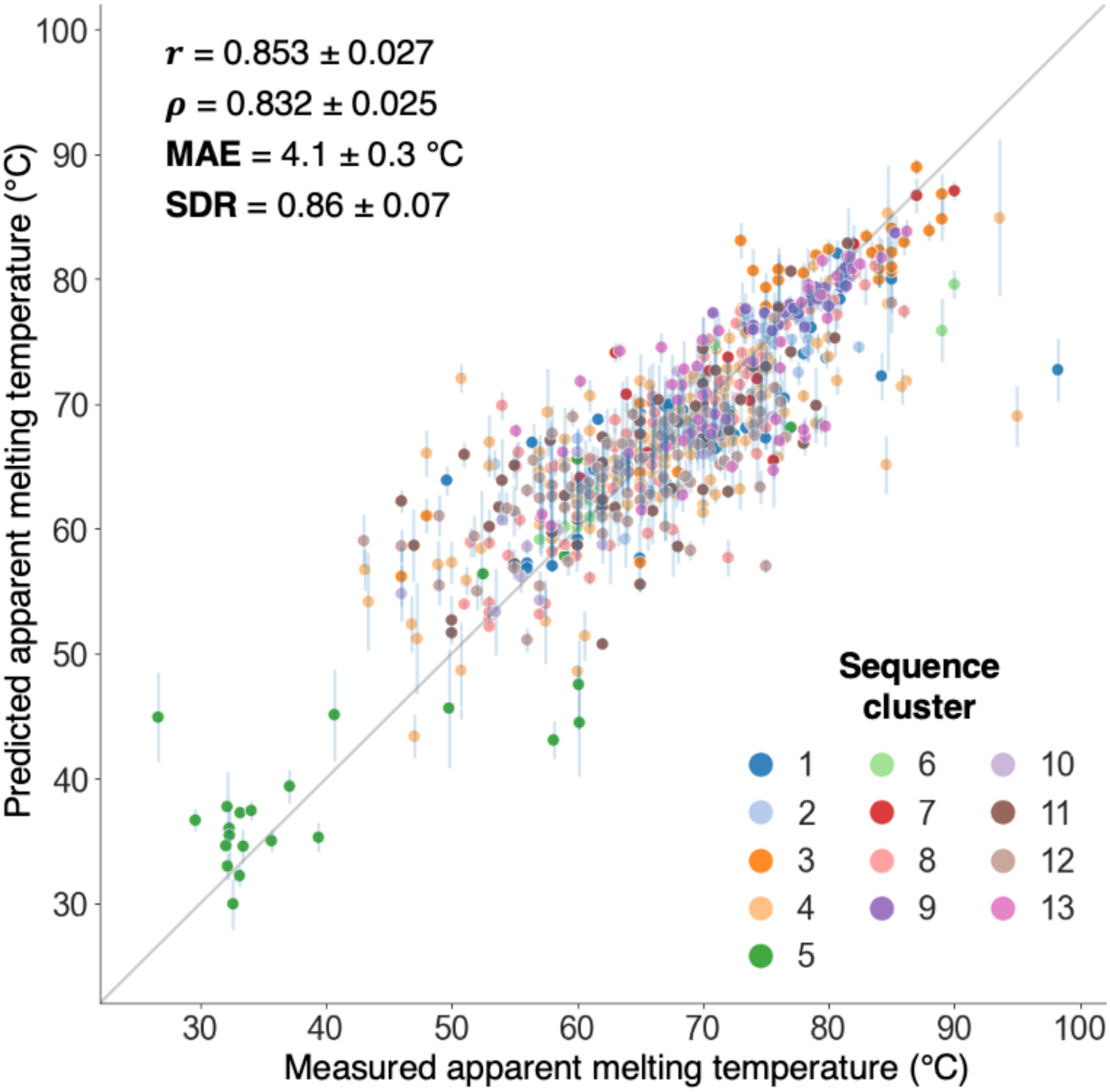
Test performance of the ensemble model across the nanobody dataset. Test-set prediction versus measured T_m_ from the final ensemble model, which consists in ridge regression stacking of models trained on diverse embeddings. For each nanobody, the prediction corresponds to the test prediction from the outer test fold of the nested cross-validation, averaged over three repeats. Error bars indicate the standard deviation of the predictions over the three pipeline repeats. The average standard deviation of the predictions over the repeats reaches 1.3 °C. Scatter point colours represent the sequence k-medoids cluster of each nanobody. The reported Pearson’s correlation (r), Spearman’s correlation (ρ), mean absolute error (MAE), and standard deviation ratio (SDR) are averaged across all outer test folds and repeats of the nested cross-validation, with corresponding standard deviations. The SDR assesses the model’s tendency to regress towards the mean value of the dataset. Overall performance metrics for the plotted data are r = 0.862, ρ = 0.845, MAE = 3.9 °C, and SDR = 0.84.

To assess model behaviour across different sequence clusters, we plotted individual cluster test-set performances (**Fig. S8**). High test performance was observed across both diverse and less diverse clusters. For instance, the highly diverse cluster 4 achieved a Pearson’s correlation of 0.741, while the less diverse cluster 9, comprising point mutations from Sulea et al. (55), reached 0.843. The consistency in test performance across clusters indicates that the model provides reliable predictions for both highly diverse nanobody sequences and point mutations of a given nanobody.

Finally, to set realistic expectations for model performance in real-world applications, we bootstrapped the Pearson’s and Spearman’s correlations calculated across subsets of our dataset, sampling from 2% to 75% of sequence predictions in test-set folds (**Fig. S9**). This analysis addresses a common misconception: when applying a predictive model to new sequences, there is often an expectation that the correlation coefficients will match those reported during the original model assessment, which is typically based on much larger test datasets. Our results show that when sampling one-third of the sequences – the proportion used in the three-fold cross-validation – the correlation coefficients exhibit a narrow normal distribution centred around r = 0.864 and ρ = 0.844, as expected (**Fig. S9**). Similar patterns are observed when sampling 10% and 75% of the sequences.

However, when evaluating performance on smaller subsets, such as a dozen sequences (2% sampling), the correlation coefficients display much broader distributions, with a 1.5 interquartile range spanning from r = 0.597 and ρ = 0.499 to nearly 1, and some outlier samples dropping almost to r = 0. These results underscore a critical point relevant to any data-driven model across various fields: reliable performance assessments require sufficiently large and diverse datasets (here approximately 50-60 sequences, as suggested in **Fig. S9**). In contrast, testing on small datasets inherently leads to substantial variability in observed performance, highlighting the unreliability of conclusions drawn from such limited assessments.

We also train an “operational” final model that we call NanoMelt, trained on the whole dataset to maximise data utilisation (see Methods), which we make available as a webserver at https://www-cohsoftware.ch.cam.ac.uk/ and for download from our repository (https://gitlab.developers.cam.ac.uk/ch/sormanni/nanomelt). NanoMelt takes around 250 sec to be run on 1,000 sequences on a single GPU (NVIDIA RTX 8000).

### 4.5 Benchmarking

We benchmarked NanoMelt against three existing methods capable of predicting melting temperatures or of providing a stability score for protein sequences (**Fig. S10**). These methods include FoldX (56), an empirical force-field-based algorithm that calculates the free energy of unfolding for a given structure; DeepSTABp (26), a transformer-based melting temperature predictor trained on over 35,000 datapoints; and sequence likelihood outputs from ESM-2 and AntiBERTy, used as proxies for stability in the context of zero-shot predictions.

Given FoldX’s reported sensitivity to minor atomistic displacements (6, 56), we selected 46 nanobodies from our dataset that have corresponding crystal structures in the PDB, to avoid biases due to inaccuracies in structural modelling. Our model achieved a Pearson’s correlation of r = 0.702 on test-set predictions on these 46 nanobodies. However, FoldX predictions showed no significant correlation with experimentally measured melting temperatures (**Fig. S10a**, r = 0.033), confirming that while empirical energy-based algorithms like FoldX are useful for predicting stability changes upon mutation, they struggle to assess stability across different but related protein sequences, such as nanobodies (57).

DeepSTABp was benchmarked on the entire dataset, but its predictions exhibited a weak Pearson’s correlation of 0.267 and a very strong regression towards the mean of its training data (**Fig. S10b**) (26).

We also tested the sequence likelihood from ESM-2 and the pseudo-log-likelihood from AntiBERTy as proxies for stability. Higher likelihood values are presumed to indicate more native-like sequences, which should correlate with greater stability, as suggested by Chungyoun et al. (58). However, these proxies did not show strong correlations with experimental data (**Fig. S10c-d**). Notably, ESM-2 achieved a Pearson’s correlation of 0.337 across the entire dataset (p-value ~ 10^−18^), outperforming stability-specific approaches like FoldX and DeepSTABp and confirming its potential for zero-shot prediction of protein fitness (3). However, further semi-supervised training or transfer learning for downstream prediction tasks appears essential to achieve accurate predictions.

### 4.6 Selection of highly stable nanobodies with NanoMelt

The most common approach to discovering new nanobodies targeting antigens of interest is through in vitro panning of VHH libraries, either obtained from immunised camelids or constructed as naive libraries. Given the short sequence of nanobodies, next-generation sequencing (NGS) has become a standard practice for gaining a comprehensive view of the nanobody repertoire post-screening. Therefore, rapid, sequence-based computational predictors of biophysical traits, like NanoMelt, can significantly enhance these discovery pipelines by enabling the selection of high-fitness nanobodies for further experimental characterisation.

To mimic this scenario and validate NanoMelt on independent data, we mined NGS-derived databases of camelid nanobody sequences (59) and selected six candidates for expression and characterisation. Selection criteria included: 1) at least 30% sequence dissimilarity from the most similar sequence in our T_m_ nanobody dataset; 2) an AbNatiV VHH-nativeness score greater than 0.85; and 3) predicted thermostability: three nanobodies should be poorly stable (predicted T_m_ < 61 °C; Nb1, Nb2, Nb3) and three highly stable (predicted T_m_ >73 °C; Nb4, Nb5, Nb6). AbNatiV (59), a recently introduced deep learning framework, accurately predicts nanobody nativeness, which relates to the likelihood of a sequence belonging to the distribution of immune-derived nanobodies. We applied a stringent nativeness filter (> 0.85) to ensure all selected sequences do not have highly unusual features that may correspond to sequencing or PCR-amplification errors, which are common in NGS datasets. The 30% minimal sequence dissimilarity to any sequence in our dataset further ensures a rigorous test of our method’s ability to generalise to sequences very distant from those it was trained on. Sequences passing the first two filters show predicted T_m_ values between 58 and 79 °C (**Fig. S11**).

We transiently transfected mammalian cells for expression (see Methods) and found that all three nanobodies predicted to be highly stable expressed well, while none of the three predicted to be poorly stable showed any expression in the supernatant (**Fig. 6** and **S12**). We then purified the three expressing nanobodies, confirmed their identity by mass spectrometry (**Table S6**), and measured their T_m_ values. Remarkably, the measured values closely matched the predicted ones, with absolute errors of 1.7, 0.6, and 0.6 °C for Nb4, Nb5, and Nb6, respectively (**Fig. 6**).

**Figure 6.**
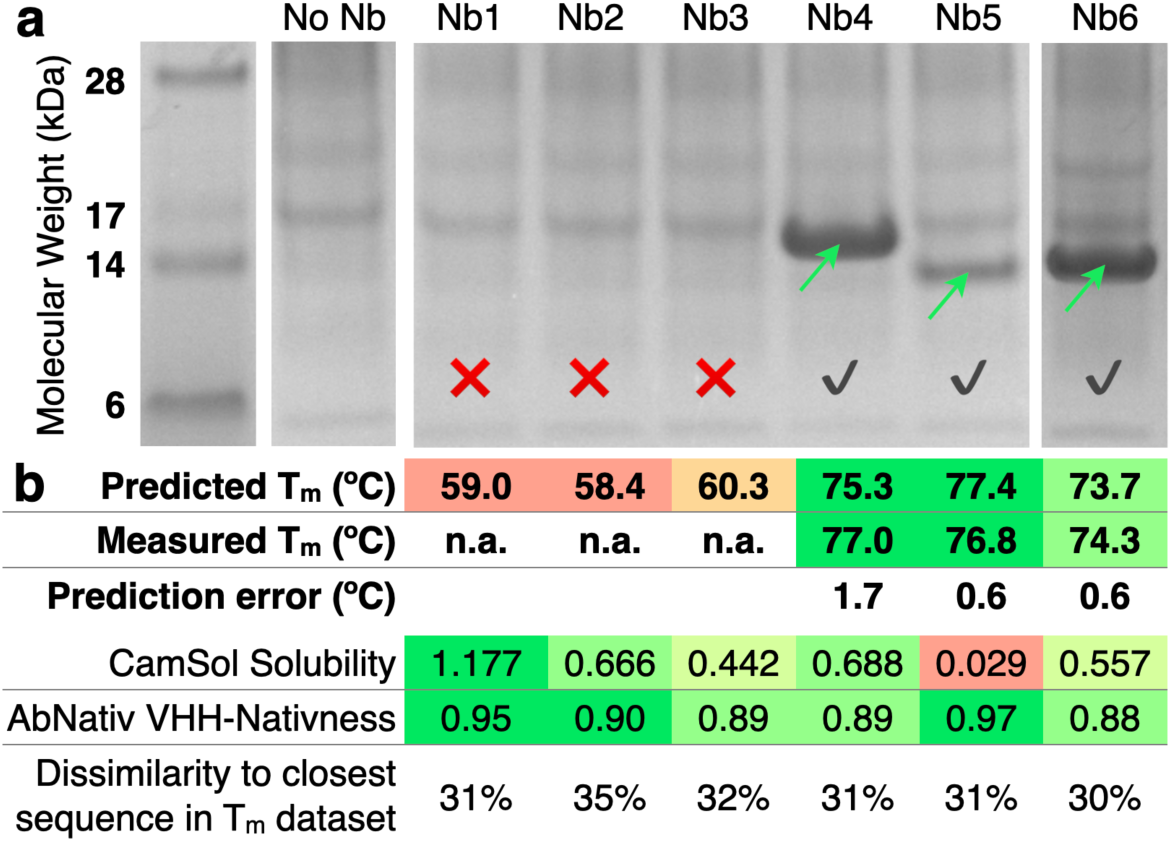
Real-world application of NanoMelt to sequences distant from those in T_m_ dataset. (**a**) SDS-PAGE analysis of mammalian cell supernatants transiently transfected with six selected nanobodies (header). The “No Nb” lane represents the supernatant of cells not overexpressing a nanobody, serving as a negative control. The Nb6 sample was run on a separate gel (see **Fig. S12** for the uncropped images and additional results from repeated independent transfections). Green arrows indicate the nanobody bands at the expected molecular weight (confirmed by Mass Spec: **Table S6**), a red cross denotes lack of expression, and a grey tick indicates successful expression. (**b**) Table summarising the predicted T_m_, measured T_m_, prediction error, CamSol intrinsic solubility score, AbNatiV VHH-nativeness, and percentage dissimilarity from the closest sequence in the T_m_ dataset. Cells related to biophysical traits are colour-coded green, yellow, and red to represent favourable, intermediate, and unfavourable traits, respectively.

It is well established that low conformational stability can impair recombinant protein expression, often leading to very low yields or complete lack of soluble expression, as observed in our case. Thus, the lack of expression aligns well with the prediction of low T_m_ values for these nanobodies. We also considered thermodynamic solubility as another factor influencing expression yield. We ran CamSol intrinsic solubility predictions, which have consistently shown very high accuracy in ranking related proteins such as nanobodies in diverse settings (8, 36, 60, 61). The results show that the non-expressing nanobodies had rather high predicted solubility, comparable to or higher than that of the two best-expressing nanobodies (Nb4 and Nb6; **Fig. 6**). Interestingly, Nb5, which has a much lower predicted solubility than all other nanobodies, also showed a visibly lower expression yield (**Fig. 6** and **S11**), supporting the idea that while conformational stability is a primary determinant of expression yield, solubility plays an important role, particularly when protein variants have similar stability.

Overall, this experimental validation on nanobody sequences that are highly divergent from those in the T_m_ dataset reinforces our confidence in the practical utility of NanoMelt. NanoMelt has the potential to streamline nanobody discovery and optimisation, offering a reliable tool for selecting highly stable, well-expressing candidates during the early stages of nanobody development.

## 5 Discussion

In this study, we conducted a comprehensive investigation into protein biophysical-trait learning using limited data and applied it to develop NanoMelt for nanobody thermostability prediction. We compiled a novel dataset of apparent melting temperatures, including 129 newly measured datapoints, resulting in a curated dataset of 640 labelled unique nanobody sequences (**Fig. 3c**). By making this dataset publicly available, we aim to provide a valuable resource for the field (**Supplementary Dataset 1**).

As is common in machine learning with limited data, our dataset is somewhat noisy, with most datapoints aggregated from the literature and obtained using varying experimental protocols and data-analysis techniques. To address this variability, we applied stratification throughout the study to ensure consistent representation of experimental methods across all training and testing splits. However, we find that explicitly adding the experimental method as input to the model does not improve performance, possibly because the biggest source of variability comes from the usage of different buffers, heating rates, and protein concentrations, an information that is not available for most entries in the dataset.

We explored multiple regression models with various sequence embeddings as input, utilising repeated nested cross-validation to prevent data leakage between model tuning and performance assessment (10). Our analysis with artificial data highlighted the importance of input representation in protein fitness prediction. The performance gap between pre-trained embeddings and categorical encodings is more pronounced with smaller datasets and narrows as the dataset size increases (**Fig. S1**).

Interestingly, embeddings pre-trained specifically on antibodies and nanobodies did not outperform those pre-trained on generic proteins, despite being reported to be better at their primary task of sequence reconstruction (39–41). For instance, ESM-derived embeddings consistently outperformed antibody- and nanobody-specific embeddings in predicting both nanobody calculated solubility scores (**Fig. S1**) and actual thermostability data (**Fig. 4a**). This finding may reflect the broader applicability of the much bigger ESM models, which are trained on larger and more diverse datasets, thus providing more comprehensive information relevant to general physicochemical properties like thermostability (62).

Notably, pre-trained embeddings also reduce overfitting, which is consistently greater at small training set sizes (**Fig. S1**) and is thus an important concern when learning from limited data, as it can hinder generalisation to unseen data (63). In this study, we systematically reported both training and testing performance to assess the tendency of various regression models to overfit. This practice, which is often overlooked in recent literature (14), ensures a transparent assessment of model performance and its likelihood to generalise to more distant sequences. To mitigate overfitting feature selection was applied to each regression model to reduce the number of free parameters. In our ensemble model, we used a diverse range of input embeddings and regressors, rather than stacking only the best-performing combinations of embedding and models, to mitigate the dependency on individual models’ biases. Moreover, only out-of-fold predictions were used as inputs when training the stacking ridge model, so that this could learn to leverage the individual predictions in a more realistic, inference-like scenario.

A key challenge with small datasets is the tendency for models to regress towards the mean, which can result in promising correlation coefficients despite a lack of coverage across the full range of measured values (64). We addressed this by calculating the standard deviation ratio (SDR), which quantifies this tendency. For example, while simple averaging of predictions performed at par or better than ridge regression stacking in terms of Spearman’s correlation, the SDR revealed a stronger tendency to regress towards the mean with averaging, leading us to prefer ridge regression for the final model (**Fig. 4b**).

While NanoMelt’s MAE is 3.9 °C (**Fig. 5**), some nanobodies exhibited much larger prediction errors, around 20 °C. These outliers often had unusually high or low melting temperatures compared to similar sequences in the training set. For instance, the W37A mutant of a nanobody with a WT melting temperature of 60.2 °C had an apparent melting temperature of 26.6 °C, but its prediction was 47.5 °C. Although the model correctly indicated strong destabilisation, it underestimated the extent of the effect, likely due to the mutation involving a hyper-conserved site.

In conclusion, we have introduced a computational pipeline that combines multiple features to enable accurate learning from small datasets (**Fig. 1**). This pipeline has the potential to be applied to the prediction of other properties and across different protein classes. Nanobodies are crucial reagents in research, diagnostics, and therapeutics, and NanoMelt can significantly streamline their development by providing a reliable and rapid tool for selecting highly stable candidates during discovery and optimisation campaigns.

## 6 Material and Methods

### 6.1 Input sequence representation

Eight distinct embeddings were studied to represent the sequences as inputs for the regression models (see **Table S1**). The categorical one-hot encoding is constructed by aligning each sequence on the 149-long AHo numbering scheme (65) via the ANARCI software (66). Each position is represented by a binary vector of size 21 filled with 0 except for a single 1 at the alphabet index of the residue under scrutiny (20 standard amino acids and a gap position). These per-position vectors are concatenated together to lead to a one-hot vector of 3,219 elements. Similarly, the physicochemical-based VHSE encoding characterises each residue with vectors of size 8 derived from the principal component analysis previously conducted on 50 hydrophobic, steric, and electrostatic descriptors (37). Gap residues in AHo-aligned sequences are represented here with 8 zeros. The per-position vectors are concatenated together to lead to a VHSE vector comprising 1,192 elements.

Two embeddings are respectively generated by the large language protein models ESM-1b (38) (650M parameters) and ESM-2 (4) (150M parameters). Both models train a transformer with an unsupervised masked language objective on sequences from a wide range of protein classes from the Universal Protein Knowledgebase (UniProt) (67). The ESM-2 model is an updated version of the ESM-1b model with a bigger training dataset (respectively, 65M and 30M sequences), and distinct architecture features (e.g., type of positional embedding). Both the ESM-1b and ESM-2 embeddings are extracted from the last layer of their transformer (respectively, the 33^rd^ and 30^th^ layers). The per-residue representations are averaged across all positions, excluding the beginning-(BOS) and ending-of-sequence (EOS) tokens, yielding 1,280- and 640-sized embeddings for ESM-1b and ESM-2 respectively.

Three additional embedding are generated by antibody-specific protein language transformers. AntiBERTy (40) (26M parameters) is a transformer trained on 42M heavy antibody sequences from the Observed Antibody Space (OAS) database (68) using a multiple instance learning framework to predict the likelihood of residues in the sequences. A per-residue representation is extracted from its last layer and averaged across all positions, excluding the BOS and EOS tokens, leading to a 512-sized embedding. The AbLang model (39) (8.5M parameters) is trained on 14M heavy antibody sequences from the OAS database with a masked language reconstruction objective. A per-residue representation is extracted from its last layer and averaged across all positions resulting in a 768-long embedding. The nanoBERT model (41) (14M parameters) follows the training protocol of AntiBERTy, but it is trained specifically on 10M nanobody sequences from the Integrated Nanobody Database for Immunoinformatics (INDI) database (69). A per-residue representation is extracted from its last layer and averaged across all positions, excluding the BOS and EOS tokens, to lead to a 320-long embedding.

Lastly, the nanobody structure predictor NanobodyBuilder2 (42) is also used to generate structure-aware embeddings. NanobodyBuilder2 is a set of four transformer models trained on ~2k nanobody structures from the structural antibody database (SAbDab) (70). A per-residue representation is extracted from the last layer of the first model (7.6M parameters) and averaged across all positions yielding a 128-long embedding.

### 6.2 Minimal dataset size search

10,000 nanobody sequences were randomly sampled from the training dataset of native camelid nanobodies of the AbNatiV nanobody nativeness model (59). Their respective structure was modelled by NanoBodyBuilder2 (42). For each nanobody, the intrinsic solubility and structurally corrected scores were computed using the CamSol algorithm (36). The intrinsic score is solely computed on the amino acid sequence, while the structurally corrected score corrects for residue solvent exposure and proximity in the modelled structure.

Both generated artificial solubility datasets were randomly partitioned with an 80:20 ratio. From the 8,000 sequences, we sampled 14 subsets of varying size ranging from 20 to 8,000 datapoints. We trained a ridge regression on each subset with hyperparameters tuned via a 5-fold cross validation by grid-search, as implemented in Scikit-learn (71). For each trained model, we computed the Spearman’s coefficient to assess the prediction performance on the held-out test set of 2,000 sequences. The process was repeated over five random splitting seeds, and the standard deviation of the performance across the five splits was plotted as error bars (**Fig. S1**).

### 6.3 Expression, purification, and preparation of nanobodies for T_m_ dataset

93 nanobody sequences were selected by cluster sampling from the Protein Data Bank (PDB)(72) to ensure diversity. The PDB was used as a source because most nanobodies therein come from different animals immunised at different time, hence providing a diverse sample, and – more importantly – because all nanobodies therein are known to have recombinantly expressed successfully and have a sequence free of any sequencing errors (which are common in NGS datasets). The selected sequences were expressed using the turbo-CHO platform (GenScript) in mammalian cells with a C-terminal His6-tag, and purified via immobilised metal affinity chromatography (IMAC) in phosphate-buffered saline (PBS) buffer. Purity was confirmed by SDS–polyacrylamide gel electrophoresis (SDS-PAGE) analysis. All protein concentrations were adjusted between 0.10 and 0.18 mg.ml^−1^ to reduce aggregation upon heating while retaining a good signal-to-noise. Nanobody concentrations were determined using blanked absorbance 280 nm values in duplicates with the Nanodrop ND-1000 instrument (Peqlab Biotechnologie, Erlangen, Germany). The extinction coefficients were calculated from the amino acid sequence using the Expasy ProtParam tool (web.expasy.org/protparam/).

### 6.4 Nano-differential scanning fluorimetry assays

The nano-differential scanning fluorimetry (nanoDSF) measurements were conducted on a Prometheus NT.48 instrument (NanoTemper Technologies, Munich, Germany) with high sensitivity capillaries. Fluorescence was monitored at 330 nm and 350 nm with an excitation wavelength of 280 nm. Temperature was increased at a rate of 2 °C/min starting from 22 °C and ending at 95 °C. All measurements were performed in duplicates in different runs using different capillaries positions to mitigate possible systematic errors.

### 6.5 Melting curve fitting

For the 191 nanoDSF curves analysed in-house, the individual fluorescence signals at 350nm and 330nm, and the 350/330 ratio trace were processed to determine a representative melting temperature per nanobody. Each fluorescence trace was first smoothed via a Savitzky-Golay filter (window length = 21, polynomial order = 2) and fitted via least-square minimisation on the following two-state thermal denaturation model (73):

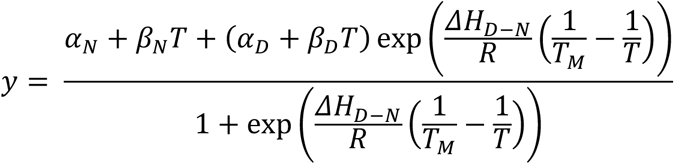

where *y* is the measured signal, α*_N_*, β*_N_* and α*_D_*, β*_D_* the intercept and slope of the linear baselines of the native and denatured states respectively, R the molar gas constant, Δ*H_D-N_* the enthalpy of equilibrium between the denatured and native states, and *T_M_* the apparent melting temperature.

A T*_M_* f 65 °C, and a ratio 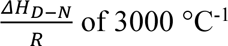 were given as initial parameter guesses. A linear fitting is done at the beginning and ending of the signal to obtain the initial guesses of the slope and intercept of the natured and denatured states. For the individual 350nm and 330nm fluorescence signals, the traces were initially corrected by subtracting it with its own fitted linear regression to provide more consistent data to the fitting algorithm thus enhancing the transition regime (a coarse but good-enough correction for the temperature-dependence of the intrinsic fluorescence). For the individual 350nm and 330nm signals, the final fitting is only carried on a window of 10°C around the *T_M_* extracted from the fitting of the 350/330 ratio trace beforehand.

For each analysed nanobody, the representative apparent melting temperature was calculated by averaging the *T_M_* values extracted from the fits of the individual 350nm and 330nm fluorescence signals, whenever conclusive. Each fit was visually inspected for accuracy; if deemed unreliable, the *T_M_* value extracted from the 350/330 ratio fit was taken instead. Among the 191 nanobodies analysed, the *T_M_* values for 166 nanobodies were extracted from the individual signals, and for 25 nanobodies from the ratio as the unfolding transition was too subtle for reliable fitting in the individual signals.

### 6.6 Construction of the nanobody thermostability dataset

A dataset of 640 nanobody melting temperatures was built in this study. The datapoints are derived from three main sources (see **Table S2**):

**–** 193 nanobody sequences from our own curated dataset, including 95 nanobodies selected from the PDB database (see **Methods 6.3**), 34 nanobodies already available in our laboratory from previous projects, and raw nanoDSF data for 64 additional nanobodies from the research conducted by Kunz et al. and published in Ref. (44). 68 nanobodies were originally published in the latter work, yet 3 could not be aligned to the AHo scheme, and the signal of NbPep53 was too poor for a T_m_ to be extracted (T_m_ likely above 90 °C with no lower plateau observed). All measurements were performed under the same experimental conditions with nanoDSF and analysed by fitting two-state thermal denaturation model (see **Methods 6.4**) to ensure high consistency. As a control, 6 nanobodies from the publication of Kunz et al. (44) were produced and characterised along our selected nanobodies. The T_m_ we obtained align closely with the data reported in the original publication of Kunz et al. (**Fig. S2**).

**–** 405 nanobodies from the NbThermo database (35). These datapoints were collated from various previously published work in the literature, and the melting temperatures were measured with a range of experimental methods including nanoDSF, DSF, circular dichroism (CD), and differential scanning calorimetry (DSC). Most of the melting temperatures reported were determined by steepest derivative analysis. The original NbThermo database of 548 nanobodies (json file downloaded in April 2023) was filtered following the alignment cleaning procedure in AbNatiV (59). In summary, some sequences were removed because:

`o` 29 entries did not have an associated sequence,
`o` 6 entries did not have a melting temperature associated,
`o` 9 sequences could not be aligned,
`o` 63 sequences matching the ones from Kunz et al. (44) were removed as we re-analysed them,
`o` 18 entries had a duplicate sequence with another entry,
`o` 18 others were duplicates of 14 sequences from our own dataset of 127 characterised nanobodies, particularly those expressed in previous project and already published.

Furthermore, 50 additional sequences have multiple melting temperatures reported for different experimental methods. The 82 unique sequences with multiple values reported (i.e., sequences duplicates or multiple experimental methods) showed a mean pairwise absolute difference of 2.4°C, which can serve as an indication of the lowest MAE that may be expected from predictors trained on this dataset (**Fig. S6**). For these sequences, the retained melting temperature corresponds to the value obtained from the experimental method when available in the following order of preference: nanoDSF, DSF, DSC, CD, and others, giving preference to our own characterisation when available.

**–** 42 nanobody sequences from two mutational studies. Specifically, 23 datapoints come from the mutational stability study comprising three WT nanobodies by Rosace et al. (8), differing by up to 4 mutations from WT. The additional 19 datapoints originate from a destabilising study on a WT nanobody biosensor in unpublished work by Dr Gabriel Ortega Quintanilla.

We make the whole dataset available in **Supplementary Dataset 1**.

### 6.7 Sequence clustering

Nanobody sequences of the curated thermostability dataset were clustered by the k-medoids algorithm from the Scikit-learn package (71). These sequences were one-hot embedded following AHo-numbering alignment prior to clustering. To determine the optimal number of clusters, the Kneedle algorithm (74) was employed to identify an elbow cutoff point. Increasing the number of clusters beyond this elbow point does not result in a significant reduction of the K-medoids’ inertia.

### 6.8 Training of the regression models

Seven regression models were tested in this study, ranging from linear models (i.e., Ridge (46), Huber (47), and Elastic Net (48)) to non-linear ones (i.e., the decision tree-based algorithms Random Forest (49) and LightGBM (50), Support Vector Machine (51), and Gaussian Process (52)). All the ranges of hyperparameter ranges given during the tuning process are directly available in the code repository at https://gitlab.developers.cam.ac.uk/ch/sormanni/nanomelt.

Each model was trained and evaluated using a repeated nested stratified cross-validation procedure (11), as illustrated in **Figure 1**. Every dataset splitting was stratified according to the type of experimental method used and the associated sequence similarity cluster. Stratification maintains the distribution of classes across the different folds during the splitting process.

The regression models were trained and ranked via a repeated nested three-fold cross-validation. This procedure incorporates two layers of cross-validation: an outer loop and an inner loop. In the outer loop, the dataset was split into three folds. Two folds were used by the inner loop to train and tune the hyperparameters on a three-fold grid search, while the remaining fold served as the test set to assess the performance of the tuned models. To enhance the reliability of the results, the entire nested procedure was repeated with three different random seeds, and the performances of each regression on the test sets were averaged. During hyperparameter tuning, each inner loop was also repeated three times, and the performances were averaged across these repeats to select the best model.

To ensure that regularisation is applied uniformly across the features (as for the Ridge, Elastic Net, SVR and GPR models), we standardized the embeddings within each cross-validation by removing the mean of the training subset and scaling it to unit variance. In addition, to reduce overfitting, we performed feature selection based on importance rankings obtained from a Ridge regression model trained on the corresponding training subset at each fold.

### 6.9 Model selection, feature selection and ensemble learning

The predicted temperatures derived from the individually trained regression models served as inputs to train the ensemble model. These predicted temperatures were collected from each outer test fold during the nested cross-validation of every model. Three sets of predicted temperatures were collected from the three repeats of the pipeline on three different seeds. An ensemble model was trained following the same stratified nested cross-validation pipeline on each set of predicted temperatures with its associated seed. The performances of the ensemble model were averaged across the three repeats.

For ensemble learning, three model selection strategies were explored to combine the various regression models. The first one selects up to the eight top ranking regression models, including the type of embedding, from the ranking of the repeated stratified nested cross-validation evaluation (**Table S3**). The second one selects the best performing regression model for each embedding type, with up to the eight embeddings (**Table 1**). And the third one selects the best performing embedding for each regression model, with up to the seven models (**Table S3**).

Three feature selection strategies were explored as inputs for the ensemble ridge model. The first one added the sequence information by concatenating the ESM-1b embedding or one-hot encoding with the eight predicted input temperatures (row A in **Table S5**). Either the size of the original embedding was maintained, or it was reduced with a linear layer to match the number of input temperatures. The second strategy added the experimental method information by concatenating it to the predicted input temperatures (row B in **Table S5**). Either the method information was added as a single integer (e.g., 0 for nanoDSF and 1 for DSF), or expanded via a linear layer to match the number of input temperatures. The third strategy integrated both sequence and experimental information into the ensemble model (row C in **Table S5**).

A ridge regression model was used as the stacking model for ensemble learning. It was trained following the same repeated stratified nested cross-validation procedure. The stacking model takes as inputs the predicted melting temperatures from the outer test folds of the previous nested cross-validation for each trained regression model.

### 6.10 Deployment of the final thermostability predictive model

For final model deployment (downloadable from https://gitlab.developers.cam.ac.uk/ch/sormanni/nanomelt, or accessible as webserver at https://www-cohsoftware.ch.cam.ac.uk/, each individual regression model was tuned first using the stratified cross-validation but then re-trained on the whole dataset. Subsequently, the stacking ridge regression and error predicting GPR were trained and re-trained following the same procedure always using as input the predicted melting temperatures from the test folds of the previous individual cross-validations.

### 6.11 Expression, purification, and characterisation of the six nanobodies of Fig. 6

Amino acid sequences encoding selected nanobodies were codon-optimized for human-cell expression using the Genscript online tool. Resulting DNA sequences were ordered from Genscript as gene fragments flanked by Golden Gate cloning sites for insertion in a pcDNA3.4 mammalian expression vector (Addgene), modified to harbour a CD33 signal secretion sequence at the N-terminus and a 6xHis tag at the C-terminus of the cloning sites. Gene fragments were cloned using Golden Gate cloning with BsmBI-v2 golden gate assembly kit (New England Biolabs; E1602S). Resulting plasmids were transformed into Dh5⍺ competent E. Coli cells (ThermoFisher Scientific) and grown overnight at 37 degrees on LB media plates containing ampicillin, before midi prep cultures were set up the next day. Midi preps were processed using QIAGEN Midi Prep kit (QIAGEN). Purified plasmids were sent for sequencing and, upon confirmation of the correct sequence, were used for transfection.

Plasmids were transfected into Expi293F cell line following protein transfection protocol from the Expi293 transfection kit manufacturer (Thermofisher Scientific; A14635). 3ml cultures were set up for each nanobody. Cells were incubated for 5 days at 37 degrees with 5% CO2 on an orbital shaker with 120 rpm. On day 5, cells were harvested by centrifugation and the supernatant was collected for SDS– polyacrylamide gel electrophoresis (SDS-PAGE) analysis and subsequent protein purification.

His Mag Sepharose Excel magnetic beads (Cytiva) were washed with PBS before being added to protein supernatants. For each 3mL culture, 100 uL of beads were added and samples were incubated on a roller at 4 degrees for 2-3 hours. Beads were then washed and resuspended with PBS to be processed on AmMag™ SA Plus Semi-automated purification System 980 (Genscript). Beads were washed with PBS at 4 mM Imidazole and protein eluted in PBS at 200mM Imidazole. Eluted proteins were then purified further by Size-Exclusion Chromatography on an AKTA system, to remove the Imidazole and to further purify the monomeric nanobodies. An AKTA pure 25 system was used with a Superdex 75 increase 10/300 GL column with PBS as a running buffer. Resulting purified proteins in PBS were aliquoted and flash frozen in liquid nitrogen. Samples were then stored in −80 °C. The mass of each protein was confirmed by comparing the theoretical molecular weight (from the the Expasy ProtParam tool) to that obtained from liquid-chromatography mass spectroscopy (LC-MS) using VION (Waters).

The apparent melting temperature of the three nanobodies that expressed was measured with a Tycho NT.6 (Nanotemper). Intrinsic fluorescence of tryptophan and tyrosine residues was recorded at 330 and 350 nm with a fixed 30 °C/min temperature ramp from 35 to 95 °C. Data were then analysed as described in the **Methods Section 6.5 “Melting curve fitting”** to obtain T_m_ values.

## Acknowledgements

We are grateful to Dr Chris Johnson for his support in facilitating the use of the Prometheus equipment at the MRC Laboratory of Molecular Biology. We acknowledge Dr Gabriel Ortega Quintanilla for sharing his thermostability data from an unpublished study. We finally thank Montader Ali, Misha Atkinson, Magdalena Nowinska, Dr Mauricio Aguilar Rangel, and Dr Oded Rimon for donating samples of their purified nanobodies to extend our dataset. P.S. is a Royal Society University Research Fellow (grant no. URF\R1\201461). We acknowledge funding from UK Research and Innovation (UKRI) Engineering and Physical Sciences Research Council (grant no. EP/X024733/1, an ERC starting grant to P.S. underwritten by UKRI).

## 7 Author Contributions

PS conceived the project with PK. PS supervised the project and secured funding. AR developed the machine learning pipeline with the help of MN. AR measured and analysed the nanoDSF curves. AR parsed and collected the nanobody thermostability data. OP and RG produced the nanobody and carried out wet-lab experiments. AR and PS wrote the first version of the paper. All authors analysed data and edited the paper.

## 8 Conflict of Interest

The authors declare no conflict of interest.

## 9 Data Availability Statement

The NanoMelt repository is accessible at https://gitlab.developers.cam.ac.uk/ch/sormanni/nanomelt. It includes all the data used in the study, the NanoMelt trained model, the python pipeline, and the notebooks used for analysis. A user-friendly webserver to run NanoMelt is provided at https://www-cohsoftware.ch.cam.ac.uk/. To access the webserver, users need to register a free account and log in

## Notes

### Competing Interest Statement

The authors have declared no competing interest.

### Summary of Updates

Consequent error on author affiliation. Needed to change the affiliation of Shimobi Onuoha to his current company.

