## Supplementary material for "Prediction of protein biophysical traits from limited data: a case study on nanobody thermostability through NanoMelt": SI

**This PDF file includes:**

- Figures S1 to S12
- Table S1 to S6
- Legend for Dataset S1
- SI References

**Other supporting materials for this manuscript include the following:**

- Dataset S1

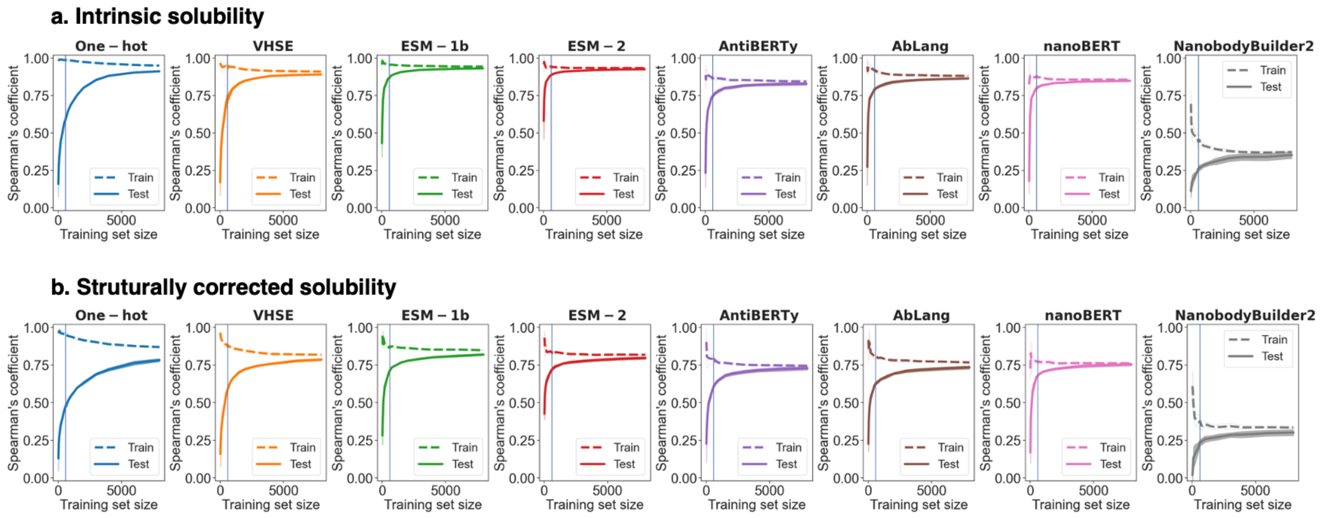


**Fig. S1. Influence of nanobody solubility dataset size on regression performance**. The intrinsic **(a)** and structurally corrected **(b)** solubility scores were computed with CamSol for 10,000 nanobodies randomly selected from the native VHH dataset of AbNatiV. 2,000 sequences were held-out for testing and 14 subsets of varying sizes, ranging from 20 to 8,000 sequences, were sampled from the remaining 8,000 datapoints. Ridge regressions were trained on each subset, representing the sequence inputs either by one-hot (in blue), VHSE (in orange), ESM-1b (in green), EMS-2 (in red), AntiBERTy (in purple), AbLang (in brown), nanoBERT (in pink) and NanobodyBuilder2 (in grey) embeddings. Performances are quantified with Spearman's coefficient (y-axis). The mean performance on training set sequences is reported in dashed line, while solid lines depict that on the 2000 test sequences in the held-out dataset. The procedure was repeated over 5 random splitting seeds, and the standard deviation of the performances is represented as shading. At 600 is drawn a vertical line.

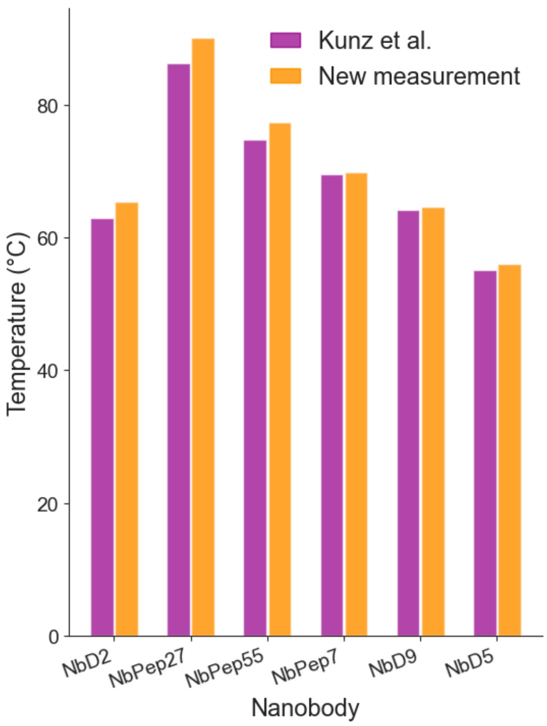


**Fig. S2. Reproducibility control between the thermostability measurements of Kunz et al. and the ones made in this study**. **(b)** Apparent melting temperatures derived from the traces measured by Kunz et al. (in purple, as reported in their publication) and by our study (in orange, fitted on a two-state thermal denaturation model, see **Methods 6.5**). The MAE between both sources is 1.8°C.

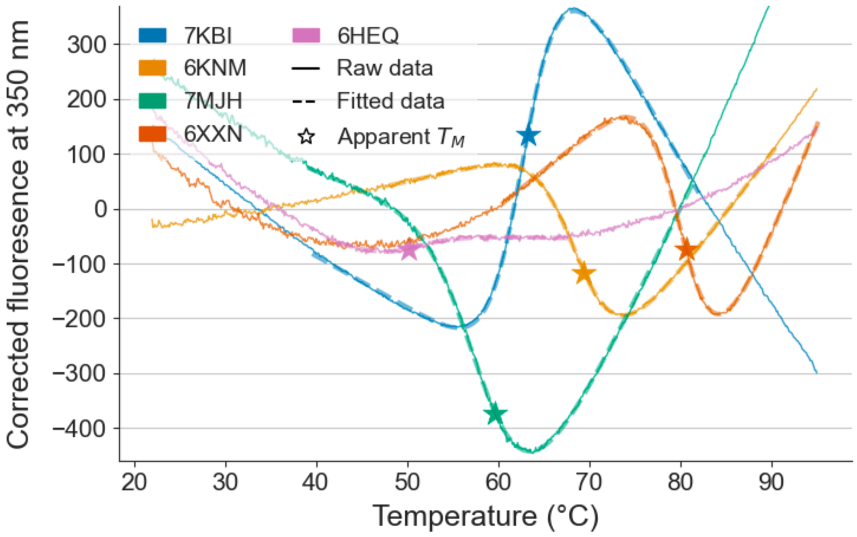


**Fig. S3. Example melting curves and fitting of five characterised nanobodies that have been selected from the PDB database**. The corrected intrinsic fluorescence at 350 nm (in solid line) was fitted onto a two-state protein denaturation model (in dashed line) to obtain the apparent melting temperature (as a star). The 350 nm traces were initially corrected for the temperature dependence of Trp intrinsic fluorescence by subtracting the measured trace with its own fitted linear regression, to enhance the transition regime and improve the quality of the fitting (see Methods). Only one replicate is shown here for clarity.

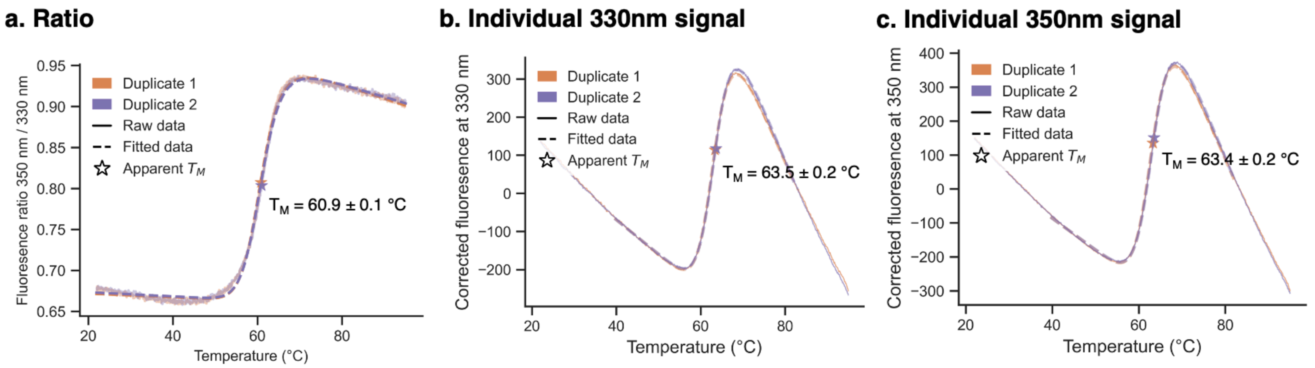


**Fig. S4. Apparent melting temperature shift of 7KBI upon ratio**. Fluorescence ratio (**a**) and individual emission at 330nm (**b**) and 350nm (**c**) from the nanoDSF signal of the 7KBI nanobody. The apparent melting temperature ($T_M$, star marker) is obtained by fitting (in dashed line) the raw data (in solid line). For the individual signals, a linear correction is applied to the signal (see Methods). The whole procedure was done in two duplicate measurements performed in separate runs (in orange and purple). The standard deviation is reported as the error on the apparent $T_M$. A shift of 3°C is observed between the apparent $T_M$s estimated from the fluorescence ratio (60.9 ± 0.1°C) and that from the individual 330nm (63.5 ± 0.2°C) and 350nm (63.4 ± 0.2°C) signals. Indeed, as reported by Žoldák et al. (1), taking the ratio of the two individual signals can introduce a bias. The two-state thermal denaturation equation for an individual intrinsic fluorescence $\boldsymbol{S}$ can be expressed as $\boldsymbol{S} = \frac{\boldsymbol{y_N}+\boldsymbol{y_D K}}{\boldsymbol{1+K}}$, with $\boldsymbol{y_N}$ and $\boldsymbol{y_D}$ the signal of the native and denatured states respectively and $\boldsymbol{K}$ an exponential term (see Methods). For the ratio of the $\boldsymbol{S_{350}}$ and $\boldsymbol{S_{330}}$ signals, this dependence becomes $\frac{\boldsymbol{S_{350}}}{\boldsymbol{S_{330}}} = \frac{\frac{\boldsymbol{y_{N350}}}{\boldsymbol{y_{N330}}} + \frac{\boldsymbol{y_{D350}}}{\boldsymbol{y_{D330}}}\boldsymbol{K}}{\boldsymbol{1} + \frac{\boldsymbol{y_{D350}}}{\boldsymbol{y_{D330}}}\boldsymbol{K}}$. The sigmoidal transition is impacted by the addition of the $\frac{\boldsymbol{y_{D350}}}{\boldsymbol{y_{D330}}}$ in the denominator, which can impact the apparent value of the $T_M$ when it differs substantially from 1. $\boldsymbol{y_{D350}}$ and $\boldsymbol{y_{D330}}$ can be fitted from their respective $\boldsymbol{y}$ signal above 85°C, assuming the denatured state is predominant. In this example, at T = 63.5°C, $\frac{\boldsymbol{y_{D350}}}{\boldsymbol{y_{D330}}}$ = 1.2, which leads to a decrease in apparent $T_M$ of a few degrees as demonstrated by Žoldák et al. (1).

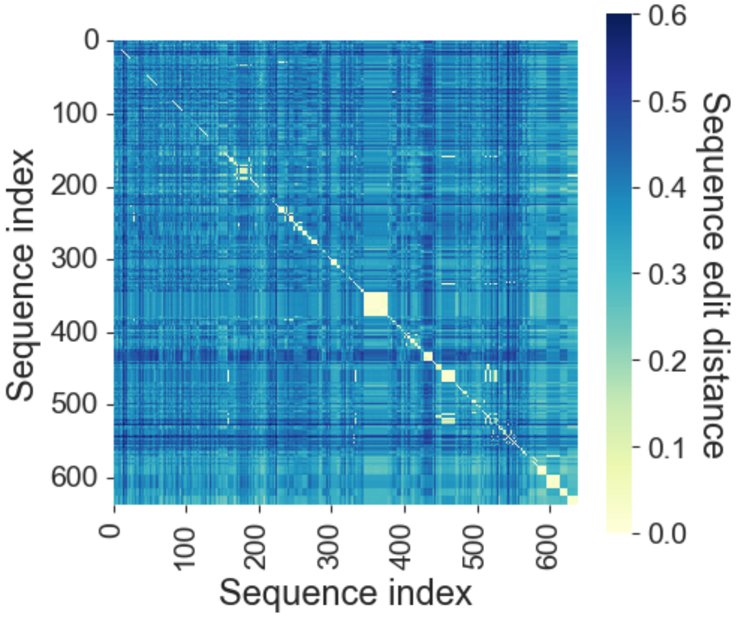


**Fig. S5. Dataset similarity heatmap**. Heatmap illustrating the sequence distances between the nanobodies in the dataset. The distance matrix is calculated based on the number of mutations (edit distance) between aligned sequences normalised by the sequence length of both sequences (see Methods).

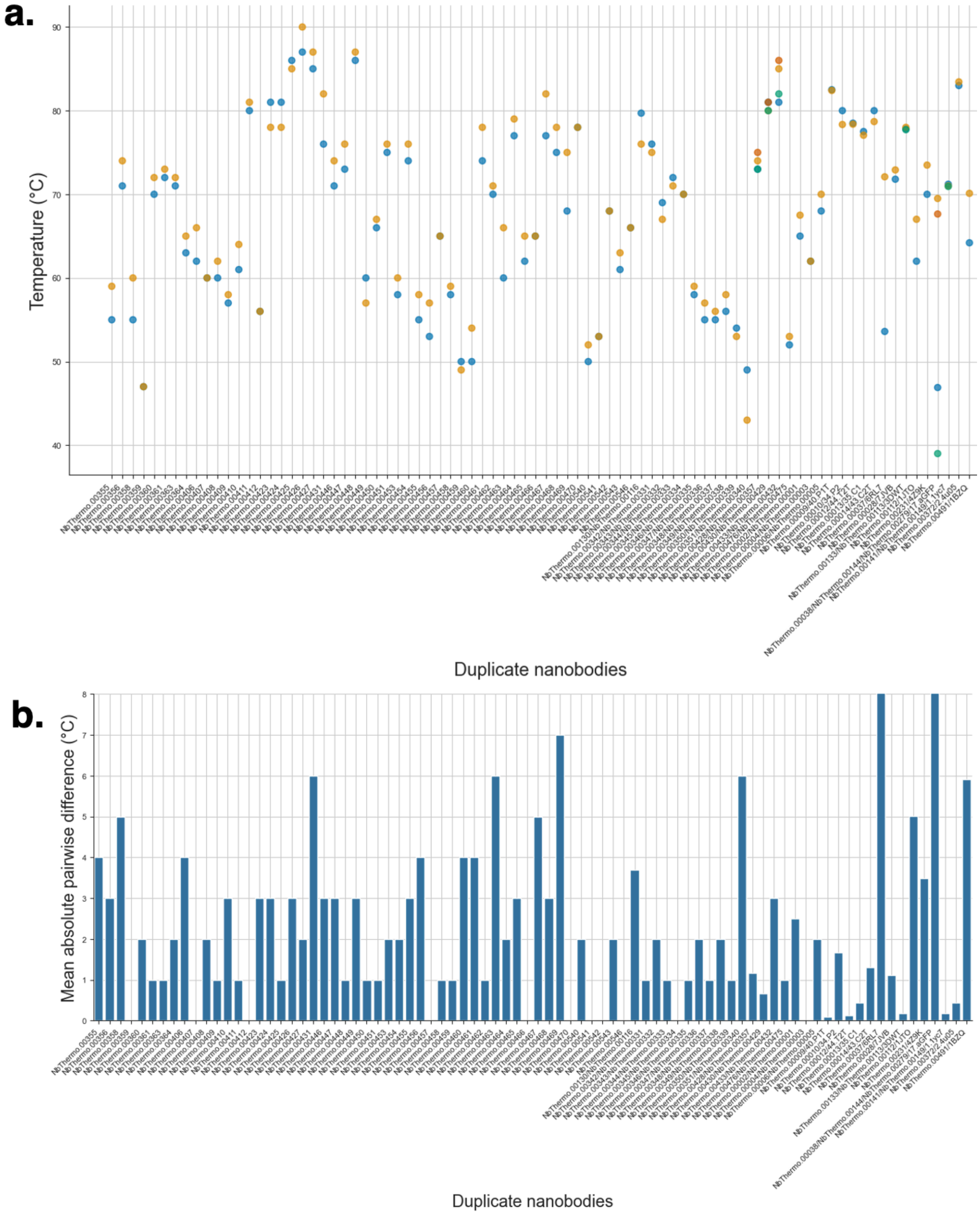


**Fig. S6. Variation in measured apparent melting temperature for the same nanobody**. **a.** Plot of the measured apparent melting temperature (y-axis) for 82 unique nanobody sequences (x-axis) for which at least two completely independent measurements were available. In this set of 82 unique nanobody sequences (x-axis), the first 50 are NbThermo (2) entries with multiple melting temperatures reported from different experimental methods, the next 18 sequences were represented by two entries, possibly reflecting different publications or the presence of different linkers/tags outside of the VHH domain, other entries were duplicates with 14 sequences from our own dataset. Colours help to distinguish the multiple values reported for each nanobody sequence. The mean pairwise absolute difference across these entries is 2.4°C. **b.** Corresponding bar plot of the mean absolute pairwise difference for each unique sequence. The y-axis is cropped at 8°C for better readability (only two outlier sequences exhibit a mean pairwise absolute difference superior to 8°C).

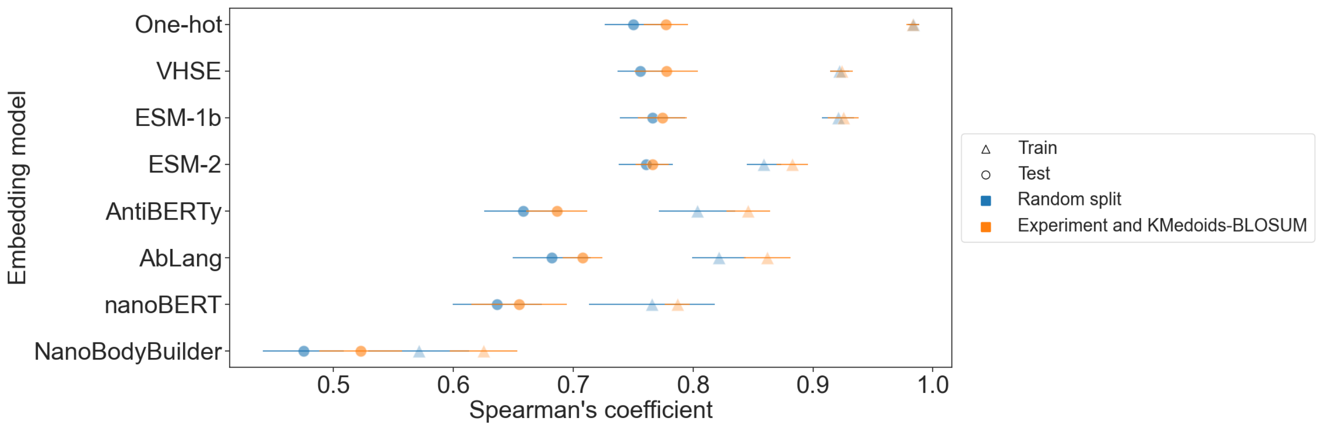


**Fig. S7. Stratification effect on regression performances.** Ridge regression trained via a nested 3-fold cross-validation. The folds were built either via random split (in blue) or via stratification on the experimental technique of melting temperature measurement and k-medoids cluster on the sequence BLOSUM identity (in orange). Round markers represent Spearman's coefficient of predictions on the test set, while triangle markers on the training set. Error bars represent the standard deviations across the cross-validation folds and repeats. With this ridge regression one-hot encoding shows among the highest test-set performance, but also the highest degree of overfitting. Conversely, ESM-2 embedding has comparable test-set performance to one-hot encoding, but substantially reduced overfitting suggesting greater ability to generalise to more distant sequences.

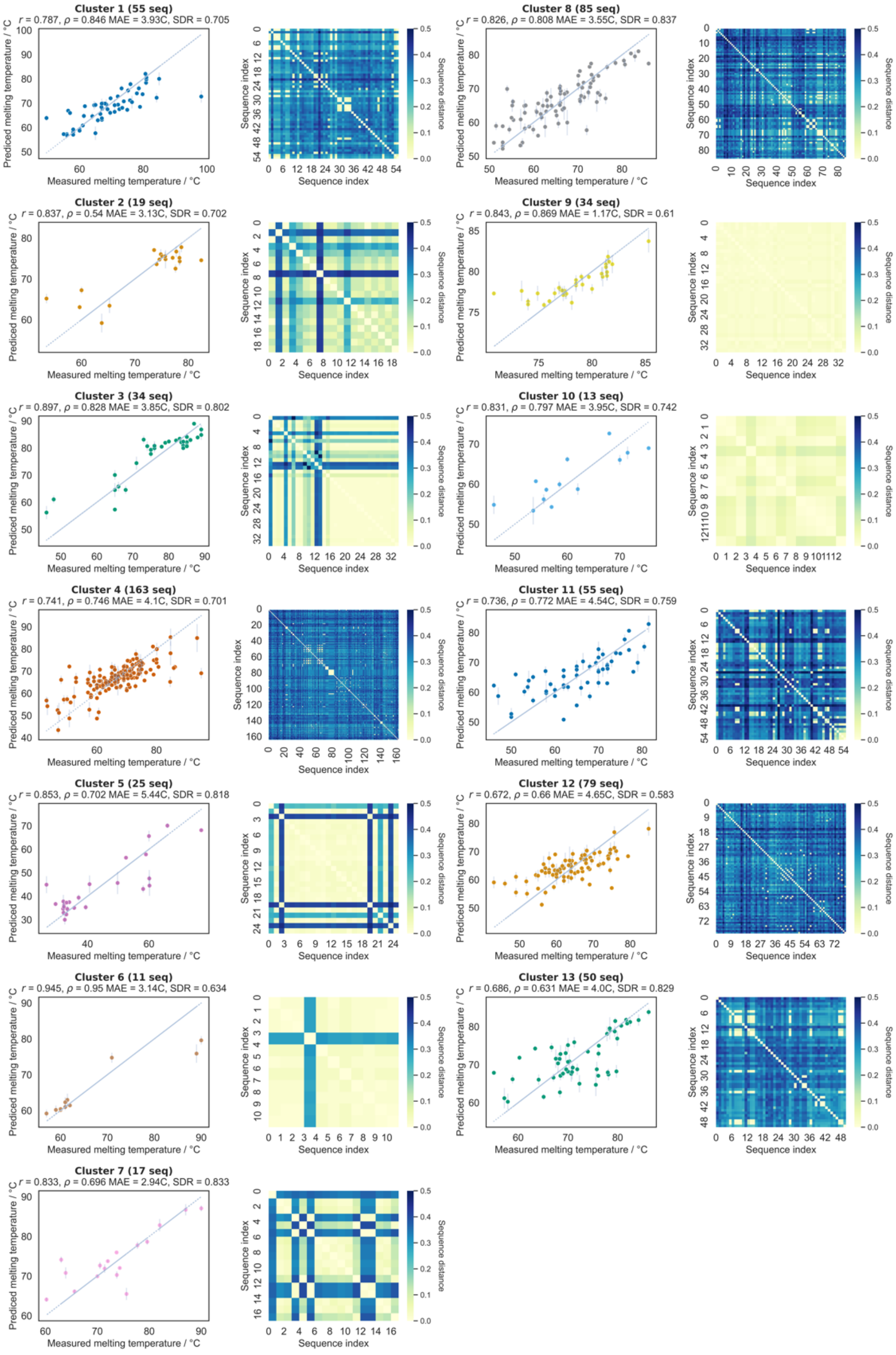


**Fig. S8. Individual cluster predictions and sequence similarity.** The test predictions are plotted for each k-medoids cluster against the corresponding measured melting temperatures (see Methods). The error bars represent the standard deviation of the predictions over the three pipeline repeats. The Pearson's coefficient (r), Spearman's coefficient (ρ), mean absolute error (RMSE) and standard deviation ration (SDR, ratio of the standard deviation of predictions over that of the measured melting temperatures) are computed within each cluster. The sequence distance matrix is calculated based on the length-normalized number of mutations between each pair of aligned sequences (see Methods).

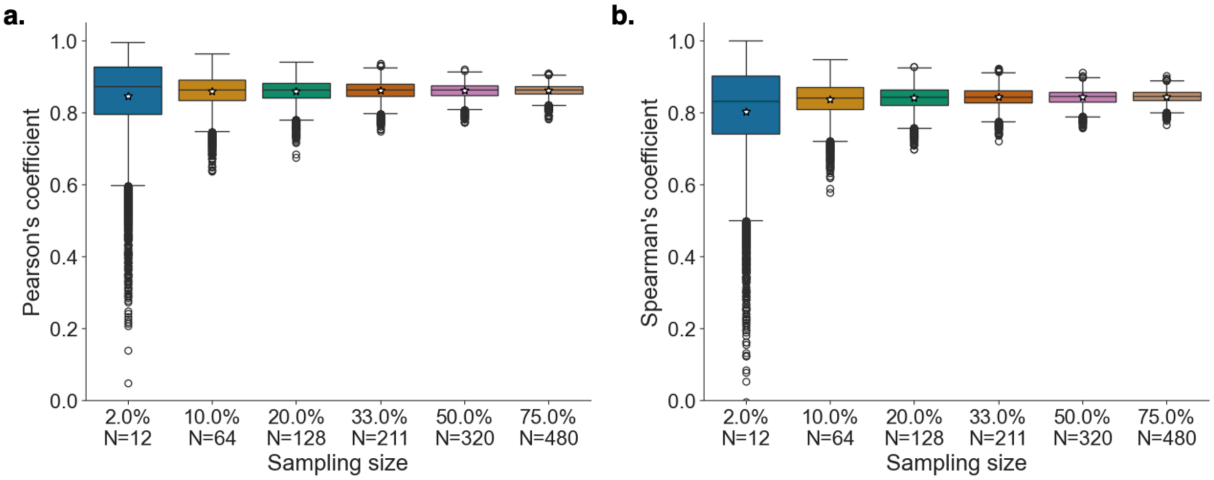


**Fig. S9. Bootstrapping of model performances** The Pearson's (**a**) and Spearman's (**b**) correlations were computed on a subset of the dataset by randomly sampling from 2% to 75% sequences from it 10,000 times. Boxes represent the first and third quartiles of the distribution, whiskers represent the 1.5 interquartile range. The median is represented as a solid line and the mean as a white star.

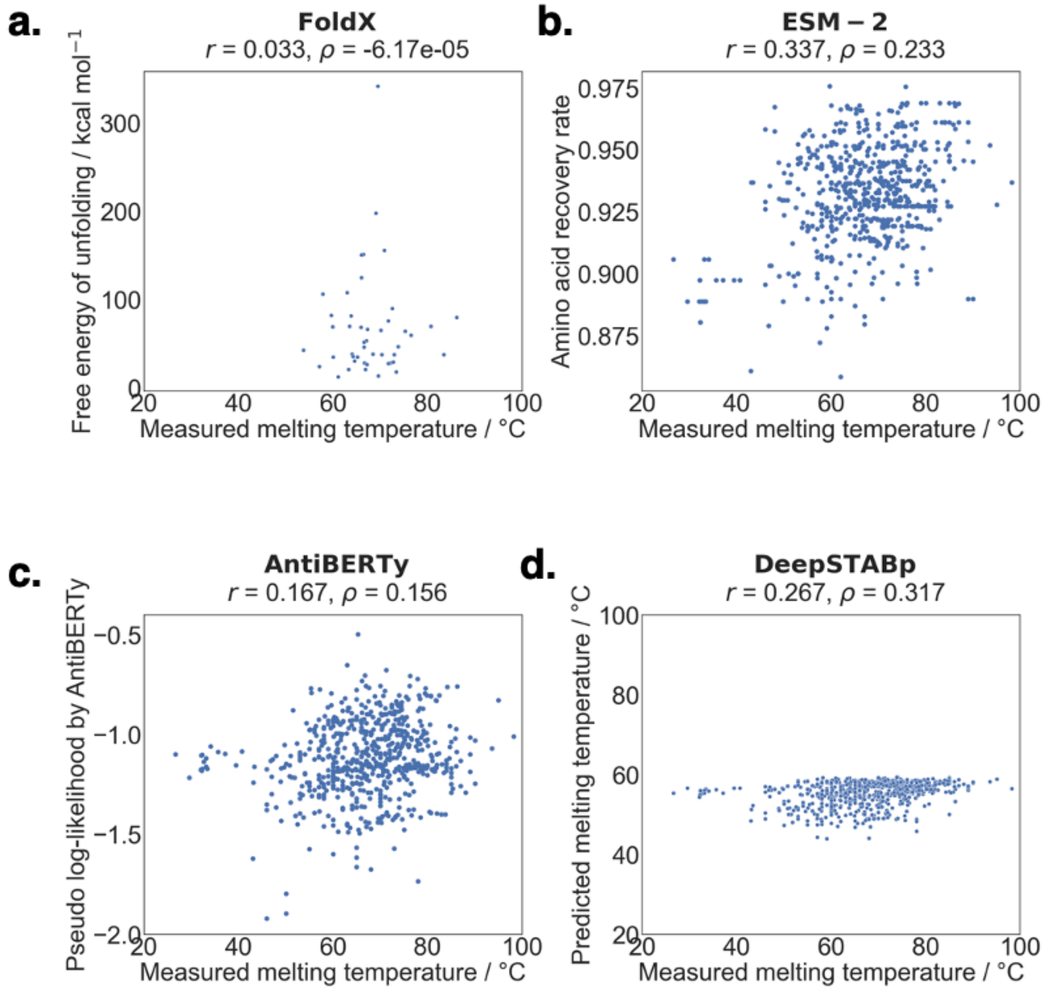


**Fig. S10. Benchmarks of existing predictive models. (a)** Predicted free energy of unfolding by FoldX for each nanobody with experimentally resolved structure in the PDB (46 out of 640) plotted with its corresponding measured $T_m$. **(b)** Amino acid recovery rate predicted by ESM-2 (masked sequence prediction) plotted against the corresponding measured $T_m$ for each of the 640 nanobodies. **(c)** Pseudo log-likelihood predicted by AntiBERTy (masked sequence prediction) plotted against the corresponding measured $T_m$ for each of the 640 nanobodies. **(d)** Predicted $T_m$ by DeepSTABp plotted against the corresponding measured $T_m$ for each of the 640 nanobodies.

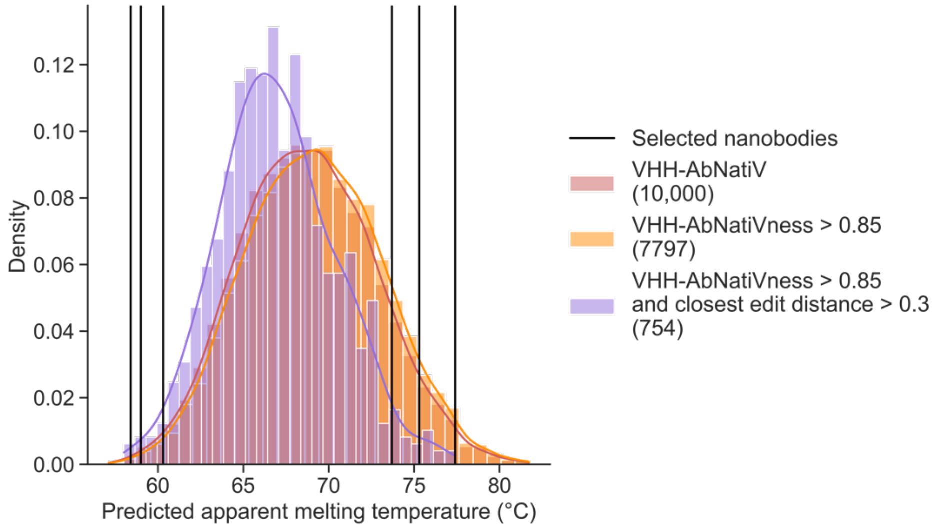


**Fig. S11. Distribution of predicted apparent $T_M$ in the AbNatiV dataset of native nanobodies.** 10,000 sequences have been randomly selected in the VHH-AbNatiV dataset of native nanobodies**(3)** and their apparent $T_M$ has been predicted with the operational predictive model we built (in red). In orange, the distribution of the predicted apparent $T_m$ for the sequences with a VHH-AbNatiV score > 0.85 among the previous sequences. In purple, the distribution of the predicted apparent $T_m$ for the sequences with at least 30% sequence dissimilarity from the most similar sequence in our $T_m$ nanobody dataset of 640 sequences. In solid black lines are represented the nanobodies selected for expression in **Section 4.6 "Selection of highly stable nanobodies with NanoMelt".**

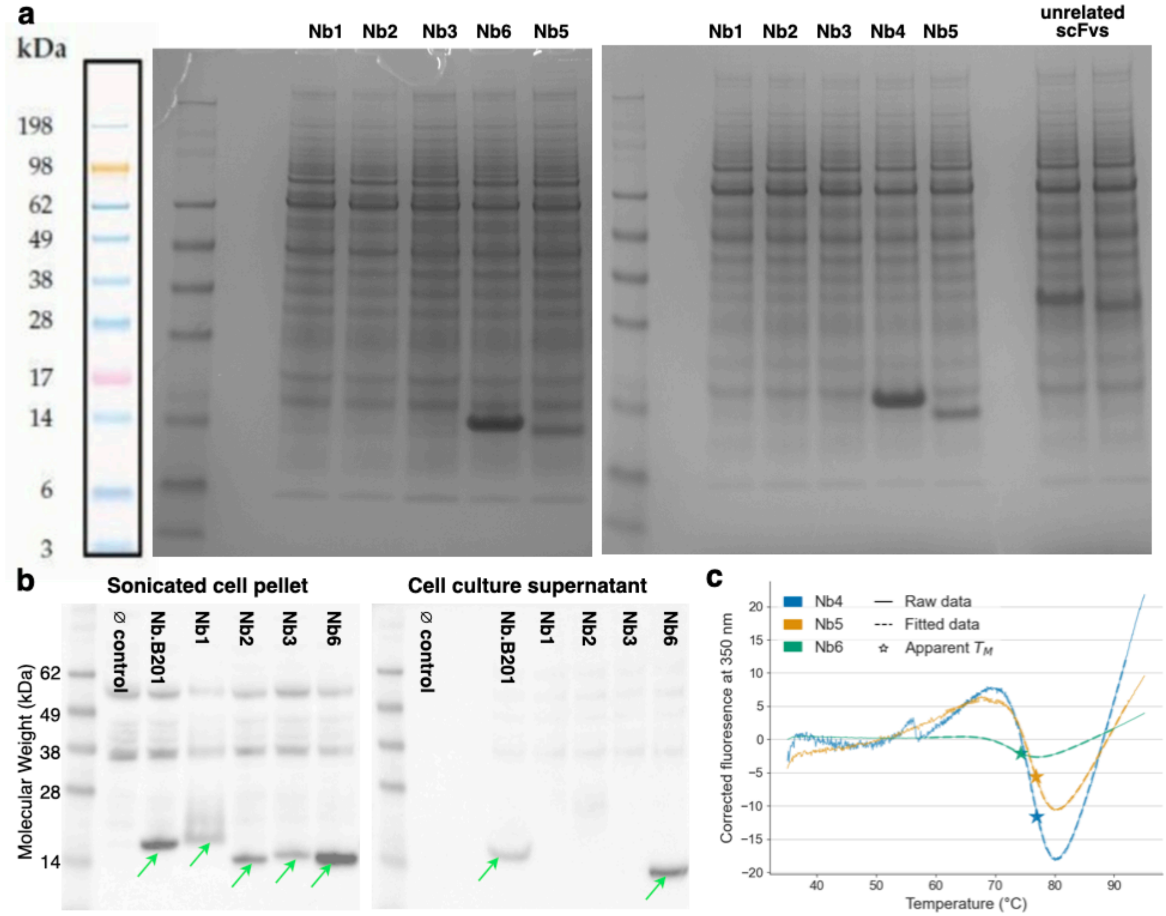


**Fig. S12. Application of the predictor to sequences distant from the $T_m$ dataset.** (**a**) Uncut SDS-page gels corresponding to the experiment shown in **Fig. 6** of the main text, with the molecular weight (MW) ladder, and the content of each lane (header). Nb6 and Nb4 expressed to very high yield, so their expression was carried out only once. Conversely, a second independent transfection and expression round was performed using a different batch of Expi293F cells to confirm the lack of expression of Nb1, Nb2 and Nb3, as well as the visibly lower expression yield of Nb5 (as inferred by the intensity of its band at around 14 kDa, compared to the bands of Nb4 and Nb6). The lanes named "unrelated scFv" (whose bands are visible at around 30 kDa) correspond to proteins being expressed at the same time, by the same cell batch, and same expression vector for an unrelated project. Here, they serve as a negative control to see what bands are found at the nanobody MW range (10-20 kDa). Such bands correspond to proteins that are normally secreted by overexpressing HEK cells during six days of over-expression. (**b**) Western blot analysis of the sonicated cell pellet (left) and the cell culture supernatant (right), from which nanobodies are usually purified. Green arrows point to the nanobody bands. Bands at higher MW are likely non-specific background signal from the anti-his-tag antibody used, as they are also visible in the empty control pellet. Nb.b201 is a nanobody that expresses rather poorly in mammalian cell, used here with Nb6, which expresses well, as a positive control. Nb1, Nb2 and Nb3 are not detected in the supernatant but present in the sonicated cell pellets, confirming their successful transient transfection. However, these samples are typically not viable for purification, as proteins found here are usually aggregated and typically still have the N-terminal signal secretion sequence attached. (**c**) The corrected intrinsic fluorescence traces of Nb4, Nb5, Nb6 at 350 nm (in solid line). The traces were fitted onto a two-state protein denaturation model (in dashed line) to obtain the apparent melting temperature (as a star). The 350 nm traces were initially corrected for the temperature dependence of Trp intrinsic fluorescence by subtracting the measured trace with its own fitted linear regression, to enhance the transition regime and improve the quality of the fitting (see Methods).

| Name | Embedding length | Number of parameters | Training dataset size | Source |
|---|---|---|---|---|
| One-hot | 3,219 | Categorical | Categorical | None |
| VHSE | 1,192 | Categorical | Categorical | Mei et al. (4) |
| ESM-1b | 1,280 | 650M | 30M | Rives et al. (5) |
| ESM-2 | 640 | 150M | 65M | Lin et al. (6) |
| AntiBERTy | 512 | 26M | 42M | Ruffolo et al. (7) |
| AbLang | 768 | 8.5M | 14M | Olsen et al. (8) |
| NanoBERT | 320 | 14M | 10M | Thorling et al. (9) |
| NanobodyBuilder2 | 128 | 7.6M | 2k | Abanades et al. (10) |

**Table S1. Sequence embeddings used in the study.** One-hot and VHSE embeddings are categorical representations. The other embeddings are latent representation derived from pre-trained machine learning models. For each embedding, the table reports the embedding length, the number of parameters and training dataset size of the trained model (where applicable), and the source. The first model of NanobodyBuilder2 is used (out of 4).

| Dataset of origin | Number of sequences kept | Content | Source |
|---|---|---|---|
| This study | 129 | 95 nanobody sequences were selected from the PDB database and expressed by Gen Script and 34 were expressed in our laboratory for previous projects. All measurements were performed under the same experimental conditions to those described by Kunz et al.. | Supplementary Dataset 1 |
| Kunz | 64 | Diverse nanobodies expressed and characterized under the same conditions. These sequences are included in NbThermo but the raw data was reanalysed in this study. | Kunz et al. (11) and Supplementary Dataset 1 |
| NbThermo | 405 | Datapoints collated from diverse sources of the literature with measurements done on a range of experimental methods. | Valdes-Tresanco et al. (2) |
| Rosace | 23 | Mutational stability study comprising three WT nanobodies differing by up to four mutations. | Rosace et al. (12) |
| Ortega | 19 | Mutational destabilising study of one WT nanobody biosensor. | Obtained upon request to Dr. Gabriel Ortega |

**Table S2. Composition of the whole nanobody thermostability dataset used in this study.** The apparent melting temperatures of nanobodies derived from three main sources: 193 nanobody sequences originate from our own curated dataset (including 129 nanobodies characterized in our laboratory and 64 in a previous work by Kunz et al (11)), 405 others originate from the NbThermo dataset (2), and 42 more from two mutational studies (23 from Rosace et al. (12) and 19 upon request to Dr. Gabriel Ortega).

| Rank | Top for each embedding | Top for each regression | Embedding | Regression | Pearson's coefficient | Spearman's coefficient | MAE (°C) | SDR |
|---|---|---|---|---|---|---|---|---|
| 1 | 1 | 1 | ESM-1b | SVR | 0.84 ± 0.028 | 0.816 ± 0.027 | 4.19 ± 0.3 | 0.87 ± 0.05 |
| 2 | | 2 | ESM-1b | GPR | 0.84 ± 0.027 | 0.814 ± 0.023 | 4.19 ± 0.27 | 0.87 ± 0.05 |
| 3 | 2 | | ESM-2 | SVR | 0.836 ± 0.025 | 0.799 ± 0.02 | 4.33 ± 0.25 | 0.87 ± 0.04 |
| 4 | 3 | | VHSE | GPR | 0.823 ± 0.028 | 0.797 ± 0.027 | 4.47 ± 0.27 | 0.84 ± 0.04 |
| 5 | | | ESM-2 | GPR | 0.831 ± 0.028 | 0.795 ± 0.022 | 4.41 ± 0.3 | 0.87 ± 0.07 |
| 6 | | | VHSE | SVR | 0.8 ± 0.037 | 0.785 ± 0.021 | 4.69 ± 0.34 | 0.82 ± 0.05 |
| 7 | 4 | | One-hot | GPR | 0.818 ± 0.031 | 0.784 ± 0.029 | 4.52 ± 0.28 | 0.82 ± 0.05 |
| 8 | | 3 | One-hot | RF | 0.815 ± 0.033 | 0.782 ± 0.027 | 4.56 ± 0.29 | 0.83 ± 0.06 |
| 9 | | 4 | One-hot | Elastic-Net | 0.823 ± 0.021 | 0.78 ± 0.018 | 4.78 ± 0.18 | 0.73 ± 0.04 |
| 10 | | | VHSE | RF | 0.814 ± 0.03 | 0.779 ± 0.022 | 4.62 ± 0.27 | 0.81 ± 0.06 |
| 11 | | 5 | VHSE | Ridge | 0.802 ± 0.031 | 0.778 ± 0.026 | 4.9 ± 0.28 | 0.77 ± 0.06 |
| 12 | | 6 | ESM-2 | LightGBM | 0.821 ± 0.02 | 0.778 ± 0.014 | 4.64 ± 0.16 | 0.82 ± 0.04 |
| 13 | | 7 | VHSE | Huber | 0.799 ± 0.033 | 0.778 ± 0.024 | 4.94 ± 0.25 | 0.77 ± 0.08 |
| 14 | | | One-hot | Huber | 0.814 ± 0.02 | 0.777 ± 0.018 | 4.69 ± 0.21 | 0.83 ± 0.06 |
| 15 | | | One-hot | Ridge | 0.815 ± 0.02 | 0.777 ± 0.018 | 4.68 ± 0.22 | 0.83 ± 0.06 |
| 16 | | | ESM-1b | Huber | 0.802 ± 0.02 | 0.775 ± 0.021 | 4.84 ± 0.26 | 0.83 ± 0.06 |
| 17 | | | VHSE | LightGBM | 0.81 ± 0.028 | 0.774 ± 0.02 | 4.7 ± 0.31 | 0.85 ± 0.06 |
| 18 | | | ESM-1b | Ridge | 0.803 ± 0.019 | 0.774 ± 0.02 | 4.84 ± 0.2 | 0.82 ± 0.04 |
| 19 | | | VHSE | Elastic-Net | 0.804 ± 0.027 | 0.773 ± 0.026 | 5.03 ± 0.28 | 0.7 ± 0.08 |
| 20 | | | One-hot | SVR | 0.807 ± 0.032 | 0.773 ± 0.032 | 4.65 ± 0.28 | 0.83 ± 0.07 |
| 21 | | | ESM-2 | Huber | 0.805 ± 0.017 | 0.767 ± 0.014 | 4.89 ± 0.2 | 0.81 ± 0.04 |
| 22 | | | ESM-2 | Ridge | 0.805 ± 0.016 | 0.766 ± 0.014 | 4.9 ± 0.15 | 0.83 ± 0.04 |
| 23 | | | ESM-1b | LightGBM | 0.805 ± 0.028 | 0.763 ± 0.024 | 4.8 ± 0.26 | 0.79 ± 0.05 |
| 24 | | | ESM-2 | RF | 0.809 ± 0.024 | 0.762 ± 0.017 | 4.82 ± 0.23 | 0.74 ± 0.03 |
| 25 | | | ESM-2 | Elastic-Net | 0.799 ± 0.018 | 0.76 ± 0.016 | 5.06 ± 0.2 | 0.74 ± 0.04 |
| 26 | 5 | | AbLang | SVR | 0.8 ± 0.026 | 0.76 ± 0.023 | 4.84 ± 0.19 | 0.89 ± 0.05 |
| 27 | | | ESM-1b | Elastic-Net | 0.786 ± 0.023 | 0.755 ± 0.017 | 5.13 ± 0.24 | 0.73 ± 0.05 |
| 28 | | | AbLang | GPR | 0.789 ± 0.021 | 0.751 ± 0.021 | 5.05 ± 0.2 | 0.83 ± 0.07 |
| 29 | 6 | | AntiBERTy | GPR | 0.786 ± 0.027 | 0.745 ± 0.036 | 5.04 ± 0.31 | 0.82 ± 0.06 |
| 30 | | | ESM-1b | RF | 0.786 ± 0.025 | 0.743 ± 0.017 | 5.08 ± 0.27 | 0.7 ± 0.06 |
| 31 | | | AntiBERTy | SVR | 0.78 ± 0.027 | 0.739 ± 0.034 | 5.01 ± 0.3 | 0.86 ± 0.06 |
| 32 | | | One-hot | LightGBM | 0.786 ± 0.029 | 0.734 ± 0.022 | 5.08 ± 0.27 | 0.82 ± 0.08 |
| 33 | | | AbLang | LightGBM | 0.777 ± 0.021 | 0.728 ± 0.017 | 5.14 ± 0.2 | 0.79 ± 0.04 |
| 34 | 7 | | nanoBERT | GPR | 0.784 ± 0.027 | 0.723 ± 0.035 | 5.11 ± 0.38 | 0.81 ± 0.06 |
| 35 | | | AntiBERTy | LightGBM | 0.771 ± 0.028 | 0.717 ± 0.032 | 5.22 ± 0.25 | 0.78 ± 0.04 |
| 36 | | | AbLang | RF | 0.764 ± 0.032 | 0.715 ± 0.029 | 5.32 ± 0.25 | 0.7 ± 0.04 |
| 37 | | | AbLang | Ridge | 0.742 ± 0.022 | 0.708 ± 0.017 | 5.64 ± 0.24 | 0.77 ± 0.05 |
| 38 | | | AbLang | Huber | 0.742 ± 0.022 | 0.704 ± 0.02 | 5.66 ± 0.25 | 0.76 ± 0.04 |
| 39 | | | AntiBERTy | RF | 0.765 ± 0.024 | 0.701 ± 0.037 | 5.34 ± 0.23 | 0.71 ± 0.05 |
| 40 | | | nanoBERT | LightGBM | 0.767 ± 0.031 | 0.694 ± 0.041 | 5.34 ± 0.26 | 0.79 ± 0.04 |
| 41 | | | nanoBERT | SVR | 0.755 ± 0.037 | 0.688 ± 0.04 | 5.4 ± 0.41 | 0.86 ± 0.06 |
| 42 | | | AntiBERTy | Ridge | 0.731 ± 0.026 | 0.686 ± 0.025 | 5.72 ± 0.24 | 0.75 ± 0.06 |
| 43 | | | nanoBERT | RF | 0.757 ± 0.039 | 0.685 ± 0.039 | 5.42 ± 0.32 | 0.7 ± 0.04 |
| 44 | | | AbLang | Elastic-Net | 0.729 ± 0.022 | 0.684 ± 0.024 | 5.82 ± 0.28 | 0.69 ± 0.04 |
| 45 | | | AntiBERTy | Huber | 0.726 ± 0.028 | 0.68 ± 0.023 | 5.8 ± 0.22 | 0.73 ± 0.09 |
| 46 | | | AntiBERTy | Elastic-Net | 0.719 ± 0.022 | 0.667 ± 0.02 | 5.92 ± 0.24 | 0.63 ± 0.04 |
| 47 | | | nanoBERT | Ridge | 0.716 ± 0.034 | 0.655 ± 0.04 | 5.89 ± 0.4 | 0.75 ± 0.04 |
| 48 | | | nanoBERT | Huber | 0.713 ± 0.037 | 0.651 ± 0.04 | 5.92 ± 0.41 | 0.74 ± 0.04 |
| 49 | 8 | | NanobodyBuilder2 | GPR | 0.744 ± 0.038 | 0.65 ± 0.045 | 5.42 ± 0.33 | 0.76 ± 0.04 |
| 50 | | | nanoBERT | Elastic-Net | 0.686 ± 0.032 | 0.629 ± 0.044 | 6.3 ± 0.42 | 0.64 ± 0.18 |
| 51 | | | NanobodyBuilder2 | RF | 0.692 ± 0.059 | 0.62 ± 0.043 | 5.89 ± 0.34 | 0.66 ± 0.04 |
| 52 | | | NanobodyBuilder2 | SVR | 0.706 ± 0.044 | 0.618 ± 0.044 | 5.7 ± 0.32 | 0.79 ± 0.04 |
| 53 | | | NanobodyBuilder2 | LightGBM | 0.716 ± 0.031 | 0.616 ± 0.027 | 5.72 ± 0.23 | 0.74 ± 0.04 |
| 54 | | | NanobodyBuilder2 | Ridge | 0.598 ± 0.041 | 0.523 ± 0.034 | 6.72 ± 0.28 | 0.63 ± 0.04 |
| 55 | | | NanobodyBuilder2 | Elastic-Net | 0.592 ± 0.038 | 0.522 ± 0.033 | 6.79 ± 0.35 | 0.64 ± 0.14 |
| 56 | | | NanobodyBuilder2 | Huber | 0.594 ± 0.037 | 0.519 ± 0.031 | 6.78 ± 0.3 | 0.62 ± 0.09 |

**Table S3. Performance of the 56 combinations of embedding and regression models.** Each of the seven regression models is trained with each of the eight types of embedding. Models are ranked on their Spearman's coefficient. The Pearson's and Spearman's coefficients and mean absolute error (MAE) were averaged on the test folds of the outer loop of the repeated nested cross-validations. The reported standard deviation is obtained across the three cross-validation repeats

| | Number of temperatures | Pearson's coefficient | | Spearman's coefficient | | MAE (°C) | |
|---|---|---|---|---|---|---|---|
| | | Train | Test | Train | Test | Train | Test |
| **a.** Top-performing models | 4 | 0.845 | 0.840 | 0.819 | 0.816 | 4.2 | 4.3 |
| | 5 | 0.851 | 0.845 | 0.827 | 0.822 | 4.1 | 4.2 |
| | 6 | 0.857 | 0.851 | 0.835 | 0.830 | **4.0** | 4.1 |
| | 7 | 0.859 | 0.854 | 0.837 | 0.834 | **4.0** | 4.1 |
| | 8 | **0.861** | 0.854 | **0.841** | 0.835 | **4.0** | 4.1 |
| **b.** Best regression model for each embedding | 4 | 0.856 | 0.853 | 0.835 | 0.832 | **4.0** | 4.1 |
| | 5 | 0.857 | 0.853 | 0.835 | 0.831 | **4.0** | 4.1 |
| | 6 | 0.859 | **0.858** | 0.839 | 0.837 | **4.0** | **4.0** |
| | 7 | 0.859 | 0.857 | 0.839 | **0.838** | **4.0** | **4.0** |
| | 8 | 0.860 | 0.857 | **0.841** | **0.838** | **4.0** | **4.0** |
| **c.** Best embedding for each regression model | 4 | 0.847 | 0.844 | 0.821 | 0.819 | 4.1 | 4.2 |
| | 5 | 0.852 | 0.848 | 0.832 | 0.828 | 4.1 | 4.2 |
| | 6 | 0.857 | 0.853 | 0.837 | 0.834 | **4.0** | 4.1 |
| | 7 | 0.855 | 0.852 | 0.835 | 0.832 | **4.0** | 4.1 |

**Table S4. Model selection study for ensemble learning.** Ensemble learning was performed with a ridge stacking model. The inputs of the ensemble model were collected from each test fold during the training and evaluation of the individual regression models. They were averaged over the three evaluation repeats. After training of the ensemble model, the Pearson's and Spearman's coefficients, mean absolute error (MAE) were averaged separately on the train and test folds of the outer loop of the repeated nested cross-validations. The number of temperature column precises the number of top models selected among the following model rankings. (**a**) The 8 best performing models (**Table S3**). (**b**) The best regression model for each of the 8 embedding types (**Table 1**). (**c**) The ensemble model was trained on each of the 7 regression models, each trained on their respective best embedding (**Table S3**). The best performance for each column is reported in bold.

| | Number of sequence features | Number of experimental features | Pearson's coefficient | Spearman's coefficient | RMSE (°C) |
|---|---|---|---|---|---|
| **a.** Baseline | 0 | 0 | **0.840** | **0.816** | **5.37** |
| **b.** Adding sequence information | 1280 (ESM-1b) | 0 | 0.828 | 0.800 | 6.09 |
| | 8 (ESM-1b) | 0 | 0.856 | 0.833 | 5.64 |
| | 3129 (One-hot) | 0 | 0.823 | 0.783 | 6.16 |
| | 8 (One-hot) | 0 | 0.858 | 0.841 | 5.60 |
| **c.** Adding experimental information | 0 | 1 | 0.862 | 0.842 | 5.52 |
| | 0 | 8 | 0.861 | 0.839 | 5.55 |
| **d.** Adding both sequence and experimental information | 8 (One-hot) | 1 | 0.858 | 0.842 | 5.60 |
| | 8 (One-hot) | 8 | 0.855 | 0.838 | 5.64 |

**Table S5. Feature augmentation study for ensemble learning.** Reported performance for augmentations of input features for the ridge stacking regression combining the best regression model of each of the 8 embeddings (**Table 1**). After training of every model, the Pearson's and Spearman's coefficients, root-mean-square error (RMSE) were averaged separately on the test folds of the outer loop of the repeated nested cross-validations. (**a**) SVR-ESM-1b baseline model, for comparison. Feature augmentation was realized concatenating: (**b**) sequence encodings (with the option of feature selection to reduce the length of sequence encodings to the same as the number of input temperatures, i.e. eight); (**c**) categorical features encoding experimental methods used for melting temperature measurements (with the option of expanding the features to a length of eight by a neural network with a hidden layer with 20 neurons); and (**d**) both sequence and experimental information as described. The best performance for each column is reported in bold.

| | Sequence | Theoretical MW | Measured MW | ΔMW |
|---|---|---|---|---|
| **Nb1** | QVQLQESGGGSVQAGGSLTLSCAASRNIFSIDAMGWFRQAPGKERELVAAMSNGG NRYYADSVKGRSTTSRDNARNTLYLQRDSLKPEDTAMYYCAASTQNSDYTPEGL QLFEYDSWGQGTQVTVSS | 14605.89 | n.a. | |
| **Nb2** | QVQLVESGGGLVQPGGSLRLSCIASGFQFSDYPMSWVRQAPGKDLEWIAQIAYDG WVSRYNDPAEGRFTTSRDNAKATLYLQLTNLKIEDTSMYYCTKDLSYRTRNWARS TRGQGTQVTVSS | 14494.09 | n.a. | |
| **Nb3** | QVQLVESGGGSVQAGGSLTLSCAVSNFPYHVYTLAWFRQAPGSQRETVATVDSNG VTKVAGSVKGRFTISRDNAKNTLYLQLNSLKTEETAMYYCSKDRRGSGPRGQGTQ VTVSS | 13091.50 | n.a. | |
| **Nb4** | QVQLVESGGGSVQAGGSLRLSCAASGSAISNLYMAWFRQAPGKEREGVAQMYSG ESSTYYADSVEGRFTITHDKARNTVYLQMNDLKPEDSAMYYCAAAVGLPDLRRQ GYLSADYYRIWGQGTQVTVSS | 14850.41 | 14831 | 19 |
| **Nb5** | HVQLVESGGGSVQAGGSLRLSCEISLYIYSSYCMGWFRQAPGKEREAVAAHYTGT AITYVADSVKGRFAISEDNAKNVLYLQMNSLQPEDTAMYYCAARQPCRLWLGYE DPGEYNIWGQGTQVTVSS | 14813.50 | 14810 | 4 |
| **Nb6** | QVQLVESGGGSVQAGGSLRLSCAASGLDIHSYCMTWFRQAPGKEREEIARIGTIAG PTYADSVKGRFTISQDKAKSIVYLQMNSLQPEDTAMYYCAADRMACLRPSVQLAA YNFWGRGTQVTVSS | 14358.18 | 14337 | 21 |

**Table S6. Sequence and MW of the six nanobodies used for further validation.** Table with the sequences of the six nanobodies used in **Fig. 6** and **S11**, and their theoretical MW (calculated using the Expasy ProtParam tool). All sequences have an additional 6xHis tag for purification at the C-terminus. LC-MS was used to measure the MW of the three nanobodies that were purified, and the observed discrepancy (ΔMW) between measured and theoretical MW is fully expected: pyroglutamination (-17 Da) is observed in HEK-cell-expressed nanobodies with an N-terminal glutamine (Q), and the presence of each disulphide bond (-2 Da) is detectable in all variants (Nb5 and Nb6 also have a second, non-canonical disulphide bond).

**Dataset S1 (separate file).** NanoMelt dataset of 640 nanobodies with their apparent melting temperature.

7. J. A. Ruffolo, J. J. Gray, J. Sulam, Deciphering antibody affinity maturation with language models and weakly supervised learning.

8. T. H. Olsen, I. H. Moal, C. M. Deane, AbLang : An antibody language model for completing antibody sequences.

9. J. Thorling, *et al.*, nanoBERT: A deep learning model for gene agnostic navigation of the nanobody mutational space. https://doi.org/10.1101/2024.01.31.578143.

10. B. Abanades, *et al.*, ImmuneBuilder: Deep-Learning models for predicting the structures of immune proteins. *Commun Biol* **6** (2023).

11. P. Kunz, *et al.*, The structural basis of nanobody unfolding reversibility and thermoresistance. *Sci. Rep.* **8**, 1–10 (2018).

12. A. Rosace, *et al.*, Automated optimisation of solubility and conformational stability of antibodies and proteins. *Nat. Commun.* **14**, 1937 (2023).
